# Neural substrates of female sexual rejection: hypothalamic pathways to the periaqueductal gray

**DOI:** 10.64898/2026.01.20.700523

**Authors:** Inês C. Dias, Nicolas Gutierrez-Castellanos, Liliana Ferreira, Ana Rasteiro, Margarida A. Duarte, Susana Q. Lima

## Abstract

Selecting an appropriate behavioral response according to one’s internal state is essential for well-being. Across the reproductive cycle, fluctuating levels of sex hormones align female behavior with reproductive capacity by modulating neuronal circuits that express hormone receptors. Sex hormone receptor-expressing neurons present along the anterior-posterior axis of the ventrolateral region of the ventromedial hypothalamus (VMHvl) are key regulators of female sexual behavior. While posterior progesterone receptor-expressing neurons of the VMHvl (pVMHvl^PR+^) are fundamental for female sexual receptivity during the receptive phase of the reproductive cycle, we have recently shown that anterior VMHvl^PR+^ (aVMHvl^PR+^) neurons are involved in rejection behavior when non-receptive. Here, we mapped the connectional architecture of aVMHvl^PR+^ neurons using viral tracing approaches. As expected, these neurons strongly project to several hypothalamic areas. Furthermore, consistent with previous reports, we show that aVMHvl^PR+^ neurons robustly project to several columns of the periaqueductal gray (PAG) along its anterior-posterior axis. Artificial activation of aVMHvl^PR+^ somas selectively recruits the dorsomedial PAG (dmPAG). Optogenetic activation of aVMHvl^PR+^ axons in the dmPAG partially recapitulates the rejection phenotype observed upon activation of aVMHvl^PR+^ somas, increasing the rate of rejections in receptive females. These findings reveal a putative pathway regulating female rejection behavior within a more complex circuit, ensuring that mating does not occur during non fertile periods.

## Introduction

Mating is costly for females, who may be harmed by seminal toxins or copulatory plugs, as well as directly through behaviors such as sexual coercion and harassment^1,2^. The physical act of mating can also cause injury^3,4^, reducing the likelihood that females will re-mate with other males. These costs can stem from sexual conflict, a phenomenon that occurs when the evolutionary interests of males and females do not fully align and the reproductive strategies that maximize fitness in one sex impose costs on the other^5^. Such conflicts can emerge before copulation as well, for example through disputes over mating rates or partner choice, or after copulation, in the form of competition over sperm utilization. In addition, mating incurs energetic and physiological costs^6^ and increases exposure to predation and disease^7,8^. Together, these factors illustrate how mating can impose significant burdens on females, thereby shaping the dynamics of sexual encounters.

For the house mouse, as in many other species, mating is restricted to occasions when all favorable conditions for successful fertilization are met, and prevented otherwise. To maximize the likelihood that copulation coincides with peak fertility, females often employ a peri-ovulation mating strategy, in which sexual behavior and receptivity are concentrated around ovulation. However, the progression to copulatory behavior ultimately depends on the integration of external cues with the female’s reproductive state, effectively forming a “permissive mating checklist”^9,10^, which in addition to female fertility, also includes sufficient food and water and reduced environmental stress^11–13^.

One of the most extensively studied regions regarding the expression of sexual receptivity and lordosis, the female receptive posture, is the ventrolateral region of the ventromedial hypothalamus (VMHvl), an area densely populated with neurons expressing sex hormone receptors^14–23^. While female receptive behavior has received great attention, the rejection behavior of non-receptive females has been largely overlooked and until recently, failure to copulate was mostly attributed to inactive VMHvl “lordosis” circuitry, which involves the posterior region of the VMHvl (pVMHvl)^20–23^. However, using naturally cycling females, our lab has recently shown the importance of active sexual rejection when females are non-receptive, and uncovered the role of Progesterone receptor-expressing (PR+) neurons in the anterior VMHvl (aVMHvl^PR+^) in the control of this behavior^24^. When non-receptive, the activity of aVMHvl^PR+^ neurons is heightened and correlated with rejection behavior in response to male’s attempts of copulation, and these neurons can bidirectionally regulate rejection behavior. The activity of this population seems to be dynamically regulated across the reproductive cycle by the amount of synaptic input it receives^24^. Still, the downstream projections of aVMHvl^PR+^ neurons and their specific contribution to the expression of sexual rejection remain unknown.

Studies using classic non-specific anterograde and retrograde tracing approaches have provided a complex map of inputs and outputs to and from hypothalamic nuclei, including the VMHvl^25,26^, as well as strong interconnectivity between them^27^. However, how the different genetically defined neural subpopulations present in the VMHvl, or the anatomical diversity across the anterior-posterior (AP) axis, fit in these complex connectivity profiles has only recently started to be unravelled^28^. Investigating the connectivity of VMHvl neurons expressing Estrogen receptor (VMHvl^Esr1+^), Lo and colleagues found that the pVMHvl^Esr1+^ subpopulation preferentially sends afferents to amygdalar/hypothalamic nuclei, while aVMHvl^Esr1+^ neurons preferentially projects to the midbrain periaqueductal gray (PAG), a premotor structure^28^.

In contrast, the characterization of the outputs of VMHvl^PR+^ neurons remains incomplete (but see Inoue et al., 2019; Yang et al., 2013)^20,28^. Although it is commonly assumed that Esr1+ and PR+ neurons are a fully overlapping ensemble, suggesting that the projections could be similar across the cells expressing these receptors, this degree of overlap is controversial. Some studies report that over 92% of PR+ neurons co-label for Esr1 in both sexes^20,29^, while others show that only ∼50% of PR+ neurons co-express Esr1, and 40-60% of Esr1+ neurons co-label for PR^30^. Whereas these differences could potentially be explained by different colocalization ratios in different species (e.g. guinea pig vs rat vs mouse), they highlight the need to better characterize the connectivity pattern of VMHvl^PR+^ neurons across the AP axis in the female mouse.

Here, we first use viral tracing methods to map the efferent connectivity of aVMHvl^PR+^ and pVMHvl^PR+^ neurons. These neurons share a high degree of output regions, however we identified anatomically and functionally distinct projection patterns, in particular to the PAG. Using optogenetics, we show that the specific pathway from aVMHvl^PR+^ neurons to the dmPAG partially recapitulates the rejection phenotype previously observed upon activation of the aVMHvl^PR+^ somas, increasing the rate of rejections in receptive females, suggesting that the aVMHvl→dmPAG pathway is involved in sexual rejection in non-receptive females.

## Results

### Projections of anterior VMHvl^PR+^ neurons

To investigate the output connectivity of the aVMHvl^PR+^ subpopulation of neurons, we unilaterally injected Cre-dependent adeno-associated vectors (AAV) carrying synaptophysin tagged with GFP (SynGFP, a synaptic vesicle protein present in axonal terminals)^31^ in the anterior VMHvl of PR-Cre female mice (Fig. 1A). A quantitative whole-brain analysis of the density of SynGFP+ puncta revealed that the terminals of aVMHvl^PR+^ neurons are present in more than 150 brain regions (Table S1), strongly projecting to hypothalamic, midbrain and thalamic regions (Fig. 1B). The output connectivity of aVMHvl^PR+^ neurons differs from that of pVMHvl^PR+^ neurons, whose terminals are mainly present in hypothalamic regions (Fig. S1A,B). Importantly, 17 out of the 161 identified outputs of the aVMHvl^PR+^ neurons are specific outputs for this subpopulation (for detail see Table S1 and Table S2). These aVMHvl^PR+^ specific targets include areas within the thalamus, such as some nuclei of the geniculate complex (IntG, IGL, SGN and LGd) and ventral posterolateral nucleus of the thalamus (VPLpc), or within the cerebral cortex, such as perirhinal area (PERI), postrhinal area (VISpor), key structures for multisensory integration^32–35^.

**Figure 1.**
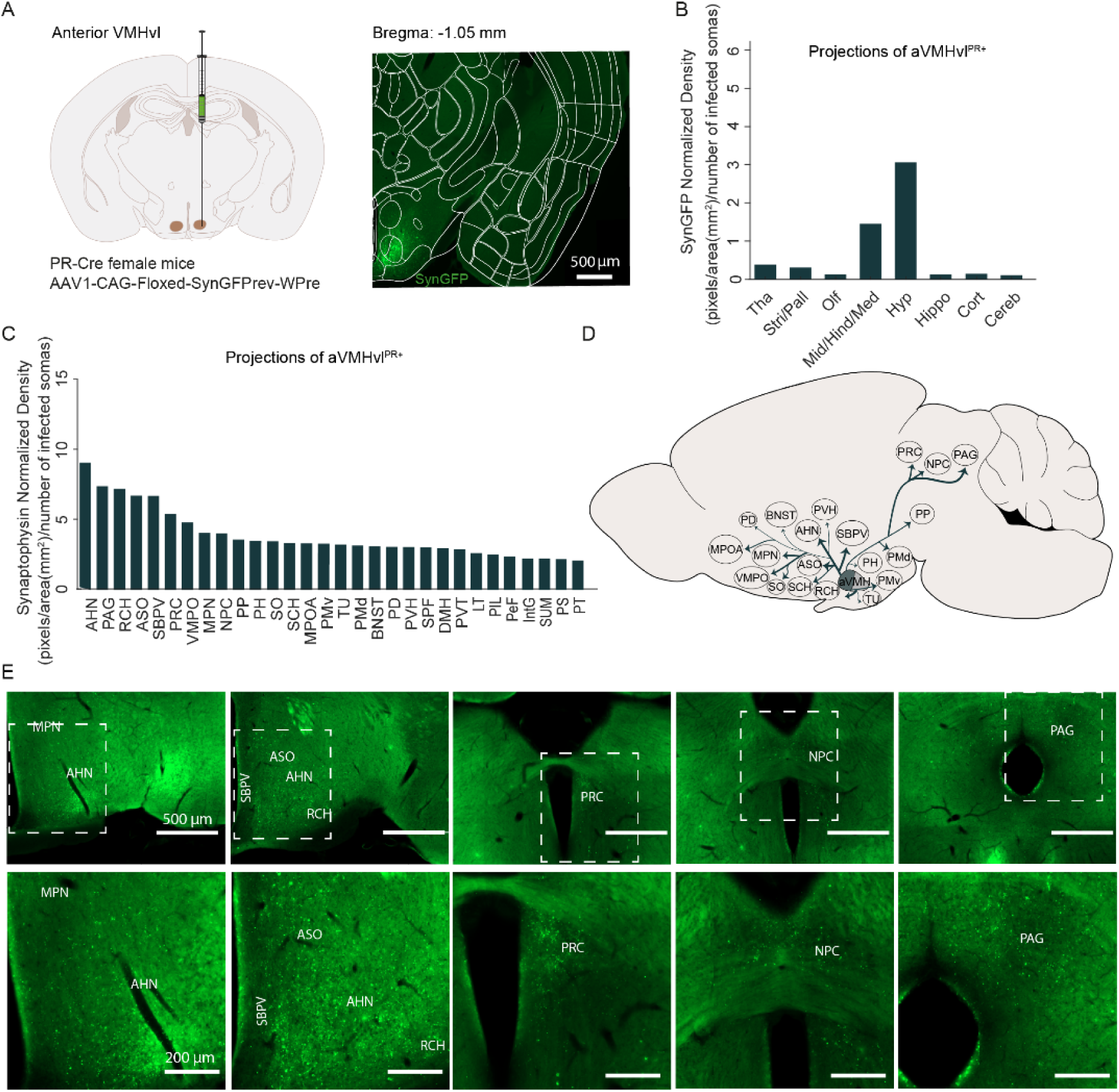
Outputs of anterior VMHvl^PR+^ neurons. (A) Schematic of virus injection aVMHvl (left) and representative histological section showing PR-Cre neurons expressing Synaptophysin-GFP (SynGFP) at the injection site (right). Scale bar: 500 μm. (B) Broad quantification of the projections of aVMHvl^PR+^ neurons to main brain divisions: thalamus (Tha), striatum/pallidus (Stri/Pall), olfactory (Olf), midbrain/hindbrain/medulla (Mid/Hind/Med), hypothalamus (Hyp), hippocampus (Hippo), cortex (Cort), cerebellum (Cereb). (C) Quantification of the projections of aVMHvl^PR+^ neurons (N = 5). The analysis of the signal across 30 principal targets revealed strong projections from the aVMHvl^PR+^ neurons to hypothalamic regions, including the anterior hypothalamic nucleus (AHN), retrochiasmatic area (RCH), accessory supraoptic group (ASO), subparaventricular zone (SBPV), medial preoptic nucleus (MPN), medial preoptic area (MPOA), ventral premammillary nucleus (PMv), dorsal premammillary nucleus (PMd); to midbrain regions, such as the periaqueductal gray (PAG), precommissural nucleus (PRC), nucleus of the posterior commissure (NPC); to thalamic regions, such as the peripeduncular nucleus (PP); and bed nucleus of the stria terminalis (BNST). (D) Schematic summary with the main output regions of the aVMHvl^PR+^. Arrow thickness represents relative projection strength. (E) Representative images of Synaptophysin-GFP signal, Scale bar: 500 μm (top) with inset on regions of interest (below). Scale bar: 200 μm.

Despite the differences in connectivity between the anterior and posterior neurons, these two subpopulations innervate many shared regions. However, within these shared regions some are anterior-preferred outputs, such as the PAG, the anterior hypothalamic nucleus (AHN), the accessory supraoptic group (ASO), the subparaventricular zone (SBPV) and the precommissural nucleus (PRC) (Fig. 1C-E, Table S1, Fig. S1E-F), while others are posterior-preferred, such as the posterodorsal preoptic nucleus (PD), the medial preoptic nucleus (MPN), the medial preoptic area (MPOA), and the ventral premammillary nucleus (PMv) (Fig. S1C-F, Table S1 and S2).

### Anterior VMHvl^PR+^ neurons innervate specific subregions of the PAG

As previously shown^28,36^, the PAG is one of the main output regions of the VMHvl, and of its PR+ subpopulation in particular^20,21^. Here, we sought to investigate in more detail the projection patterns of aVMHvl^PR+^ within the PAG (Fig. 2A,B). To do so, we quantified the synaptophysin-GFP density along the AP axis of the PAG after specifically transfecting aVMHvl^PR+^ neurons (Fig. 2C,D).

**Figure 2.**
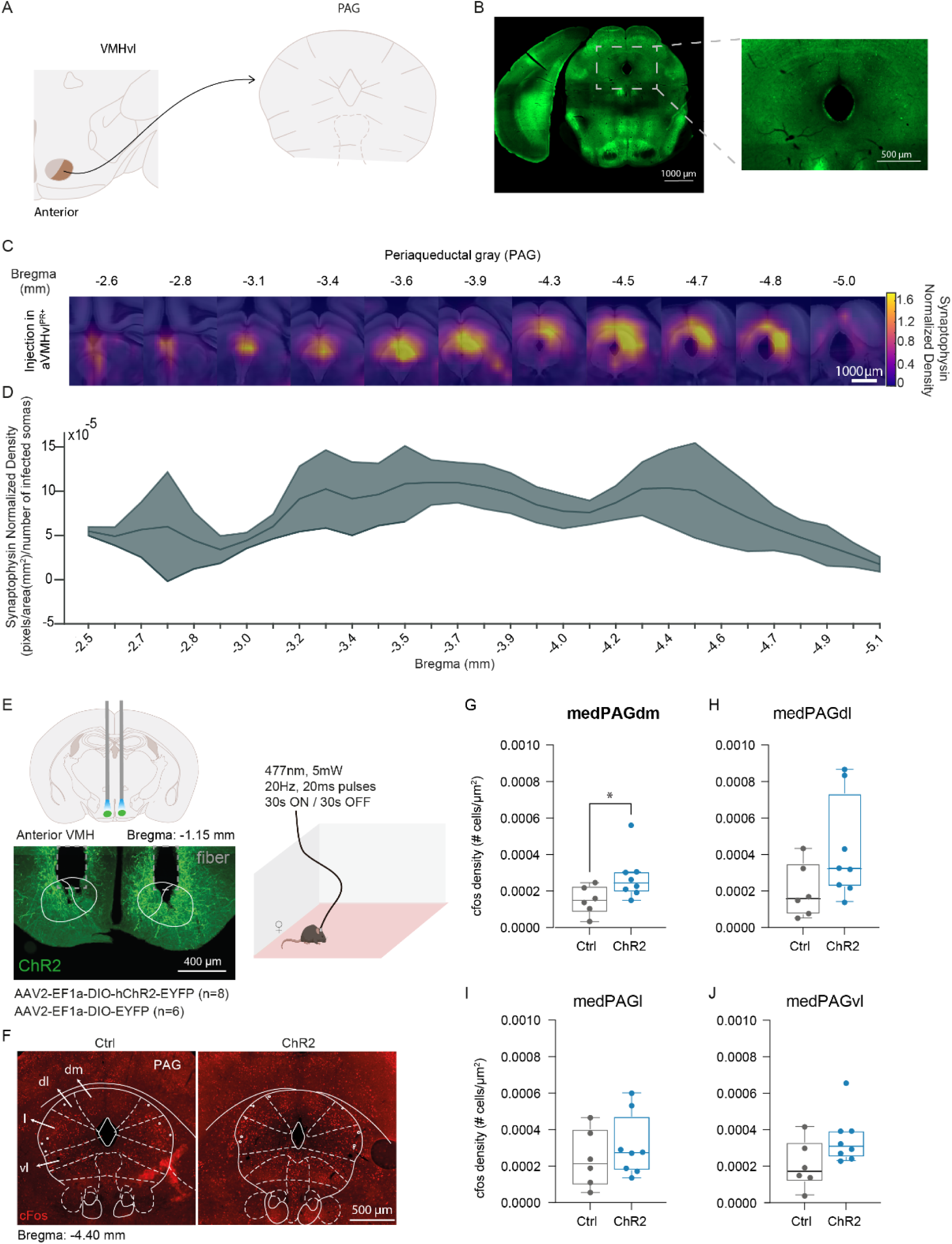
Optogenetic activation of aVMHvl^PR+^ neurons elicited an increase in cFos expression in the medial region in the AP axis of the dorsomedial column of the PAG (dmPAG). (A) Schematic of the quantified projections from the aVMHvl^PR+^ neurons to the PAG. (B) Representative image showing SynGFP expression in the PAG. Scale bar: 1000 μm (inset: scale bar 250 μm). (C) Heatmaps of binned SynGFP signal across different AP levels, normalized to injection size and averaged across mice (N=5). Scale bar: 1000 μm. (D) Quantification of aVMHvl^PR+^ to PAG projections shown in C. (E) Schematic representation of the optogenetic activation of aVMHvl^PR+^ neurons (top left), an example histological image of the injection site and fiber placement (bottom left), and the light stimulation protocol (right). Scale bar: 400 μm. (F) Representative histological images showing a cFos increase in the dmPAG of ChR2 females compared to the control group. Scale bar: 500 μm. (G-J) cFos density in the different columns of the medial PAG. (G) Mann-Whitney test, p = 0.04 and (J-L) posterior. *p < 0.05, Mann-Whitney test. Gray: Ctrl, N = 6, blue: ChR2, N = 8.

We observed that aVMHvl^PR+^ neurons project to the entire AP extent of the PAG, with stronger density in its medial and posterior portions (Fig. 2C,D). When investigating each AP level in detail, we observe a high density of aVMHvl^PR+^ projections to the dmPAG, dlPAG and lPAG columns in the dorso-ventral and medio-lateral axis (Fig. 2C). Interestingly, we observed that the projection pattern of aVMHvl^PR+^ neurons differs from that of pVMHvl^PR+^ neurons, since the posterior population projects mainly to the most posterior region of the PAG (Fig. S2A-D), preferentially targeting the lPAG and the vlPAG (Fig. S2C).

### Activation of anterior VMHvl^PR+^ somas lead to localized activation of the PAG

To determine if and how the anatomical connectivity between the aVMHvl^PR+^ neurons translates into the activation of the PAG, we optogenetically activated the somas of aVMHvl^PR+^ neurons *in vivo* and used immunohistochemistry to detect the presence of the cFos protein, a proxy for neuronal activation, in brain sections spanning the whole AP axis of the PAG (Fig. 2E).

We observed that the artificial activation of aVMHvl^PR+^ neurons induces a visible increase in cFos positive somas across the AP axis of most PAG columns. However, the cFos increase on the medial part of the dmPAG was particularly high, reaching statistical significance when compared to control animals (Fig. 2F-J, Fig. S2E-L) and suggesting that this region of the PAG might be involved in sexual rejection behavior.

### *In vivo* optogenetic manipulation of anterior VMHvl^PR+^ axon terminals in the dmPAG elicits rejection behavior in sexually receptive females

To investigate the role of the aVMHvl^PR+^ - dmPAG pathway in female sexual rejection behavior, we optogenetically activated the axon terminals of aVMHvl^PR+^ neurons projecting to the dmPAG of naturally cycling, sexually receptive females (proestrous-estrous, PE) during a sexual encounter. To do so, adult PR-Cre females were bilaterally injected in the aVMHvl with a viral construct carrying either ChR2 (ChR2) or YFP (Ctrl), and implanted with a single fiber-optic cannula above the dmPAG (Fig. 3A, Fig. S3A). After at least 3 weeks of recovery, PE females were allowed to interact with a stud male. The stimulation pattern consisted of cycles of 1 s light ON (20 Hz, 5 ms, 3-4 mW) and 4 s light OFF (stimulation protocol based on Ahmadlou et al., 2024; Cao et al., 2025, see Methods for details, Fig. 3B)^37,38^. To assess the efficiency of our stimulation protocol, we unilaterally injected a viral construct expressing ChR2 in the aVMHvl of PR-Cre females and performed *ex vivo* whole-cell patch clamp recordings in brain slices containing the PAG (Fig. 3C). The stimulation reliably evoked EPSCs in PAG neurons with high fidelity *ex vivo* (Fig. 3D).

**Figure 3.**
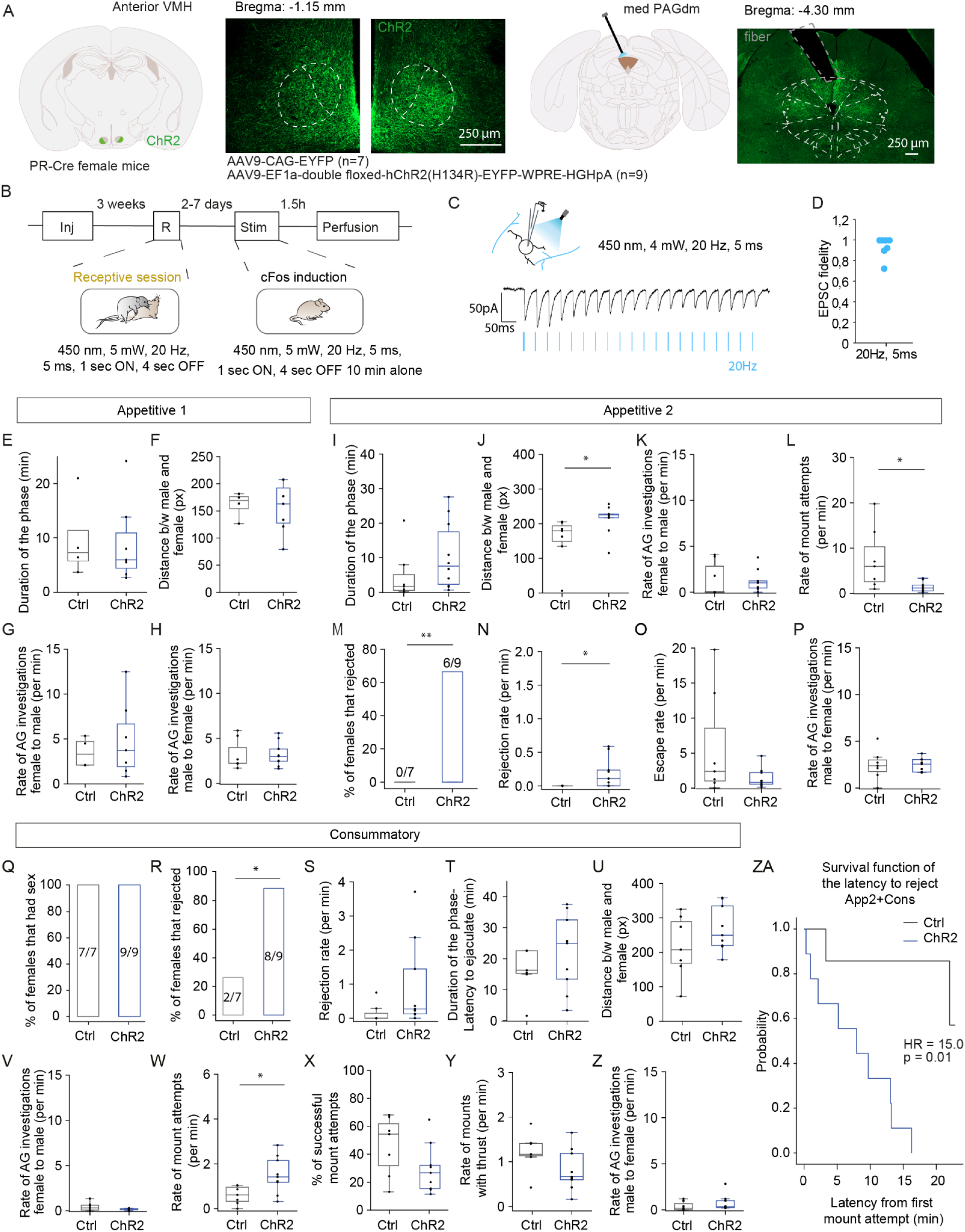
Optogenetic activation of the aVMHvl^PR+^-dmPAG pathway leads to an increase in rejection behavior during App2. (A) Schematic representations and representative histological sections of the ChR2 virus injection in the aVMHvl (left) and the mono fiber-optic cannula implantation in the dmPAG (dashed lines represent optical fiber placement). Scale bars: 250 µm. (B) Behavioral design (see Methods). (C) Schematic representation of whole-cell recording of PAG neurons while stimulating aVMHvl^PR+^ axons expressing ChR2 (top) with an example of evoked EPSCs with a 20 Hz, 5 ms, 4 mW light stimulation protocol (bottom). (D) EPSC fidelity (EPSC per light pulse) during the light stimulation protocol (n = 7 neurons from N = 4 mice). (E) Duration of Appetitive phase 1 (App1), equivalent to the latency to the first mount attempt by the male from the moment he enters the experimental arena. (F) Average distance between the couple. (G) Rate of male-directed anogenital investigations by the female during App1. (H) Rate of female-directed anogenital investigations by the male. (I) Duration of Appetitive phase 2 (App2), from the first mount attempt to the first successful mount by the male. (J) Average distance between the couple during App2. Mann–Whitney U test, p = 0.023. (K) Rate of male-directed anogenital investigations displayed by females during the App2 phase. (L) Rate of mount attempts performed by the male. Unpaired t-test, p = 0.025. (M) Percentage of females that displayed at least one rejection event during App2. N-1 Two-proportion test, two-tailed p = 0.008. (N) Rate of rejection events during App2. Mann–Whitney U test, p = 0.015. (O) Rate of escapes performed by the female during App2. (P) Rate of male-directed anogenital investigations by the female during App2. (Q) Percentage of females that had sex. (R) Percentage of females that displayed at least one rejection event during Consummatory phase (Cons). N-1 Two-proportion test, two-tailed, p = 0.017. (S) Rate of rejection events during Cons. Mann–Whitney U test, p = 0.052. (T) Duration of the Cons phase, from the first successful mount by the male to ejaculation or session termination. (U) Average distance between the couple during Cons. (V) Rate of male-directed anogenital investigations displayed by females during the Cons phase. (W) Rate of mount attempts performed by the male. Unpaired t-test, p = 0.018. (X) Percentage of successful mounts during Cons. (Y) Rate of mount with thrusts (successful mounts) performed by the male. (Z) Rate of male-directed anogenital investigations by the female during Cons. (ZA) Survival analysis of the latency to the first rejection. ChR2 females showed a significantly higher probability of rejecting earlier than controls (Cox proportional hazards model, HR = 15.0, 95% CI = 1.8–123.7, p = 0.01). *p < 0.05, **p < 0.01. Boxplots represent the median and interquartile range (IQR); whiskers extend to 1.53 IQR. Gray: Ctrl N = 7, blue: ChR2 N = 9.

Contrary to the observed when optogenetically activating aVMHvl^PR+^ somas^24^, activation of dmPAG terminals did not alter the behavior of test females during the Appetitive 1 phase of the social interaction (from male in until the first male mount attempt), when compared to Ctrl animals (Fig. 3E-G). Males in both groups also investigated the female at similar rates during the Appetitive phase 1 (Fig. 3H).

The duration of the Appetitive phase 2 (from the first male mount attempt until the first successful penile intromission) for ChR2 females showed a slight, not significant (Fig. 3I) increase compared to controls, and was not correlated with the number of rejections (Fig. S3). The distance between the couple in the ChR2 group during this phase was significantly increased (Fig. 3J), with no difference in the rate of male-directed anogenital investigations by the females (Fig. 3K). The rate of male mount attempts was significantly decreased in the ChR2 couples (Fig. 3L). These differences may arise from the fact that a significantly higher proportion of ChR2 females (ChR2: 6 out of 9) displayed sexual rejection (boxing and kicking) compared to Ctrl females (Ctrl 0 out of 7) (Fig. 3M), with a significantly higher rejection rate as well (Fig. 3N). No difference was observed in the rate of escapes performed by the females of the two groups (Fig. 3O). During the Appetitive phase 2, males in both groups also investigated the female at similar rates (Fig. 3P), suggesting that the disruption in the social interaction is not caused by a decrease in motivation from the males interacting with the ChR2 females.

Although the artificial activation of the aVMHvl^PR+^ - dmPAG pathway is sufficient to shift the behavior of PE females towards sexual rejection, the manipulation was not sufficient to prevent copulation (Fig. 3Q). All females, independently of displaying rejections during the Appetitive phase 2 or not, accepted mounting by the male and allowed him to ejaculate. However, when we examined the behavior during the Consummatory phase (from the first successful penile intromission until the male ejaculates), we observed differences between the behavior of the two groups. A larger percentage of ChR2 females performed rejection behavior during the Consummatory phase, compared to the Ctrl females - 8 out of 9 ChR2 females displayed rejections, while only 2 out 7 Ctrl rejected (Fig. 3R,S). Even though more ChR2 females rejected, the rejection rate between the two groups was not different (Fig. 3S). The duration of the phase also did not differ between the two groups (Fig. 3T), nor the distance (Fig. 3U). The rate of male-directed anogenital investigations performed by the female also did not differ (Fig. 3V). However, in the ChR2 couples, the males performed a significantly higher rate of mount attempts (Fig. 3W), even though a large percentage of those mounts attempts did not end in penetration (Fig. 3X). The rate of successful mounts was not altered (Fig. 3Y), nor did the rate of anogenital investigations performed by the male (Fig. 3Z).

Interestingly, we observed a positive correlation between the number of rejections during the Consummatory phase and the duration of the phase (Fig. S3C), but no relationship between the number of rejections during the Appetitive phase 2 and the duration of the Consummatory phase (Fig. S3D). During the Consummatory phase, the number of mount attempts performed by the male is positively correlated with the number of rejections performed by the ChR2 females (Fig. S3E), suggesting that rejection behavior leads to an increase in motivation of the male to mate (Fig. 3W). Finally, a closer look at the pattern of mount attempts and rejections showed that the Ctrl females rejected towards the end of the session, while the ChR2 females rejected earlier, after the first mount attempt performed by the male (Fig. S3F). This observation is supported by survival analysis of the latency to the first rejection - ChR2 females exhibited a significantly higher probability of rejecting than Ctrl (∼15-fold higher hazard), shifted towards earlier time points (Fig. 3ZA).

The aVMHvl^PR+^ - medial dmPAG terminal stimulation did not have an effect in other behavioral metrics, such as total distance traveled, speed and center occupancy when the female was alone in the cage (Fig. S3G,H), suggesting it does not affect locomotion and it is not anxiogenic; importantly, the stimulation of ChR2 females did not affect social encounters with female conspecifics (Fig. S3I), supporting the specificity of the manipulation to the mating context. In addition, activation of aVMHvl^PR+^ terminals in the dmPAG did not increase cFos in aVMHvl^PR+^ neurons, suggesting that the change in behavior is unlikely to be due to the recruitment of other output regions of the aVMHvl^PR+^ neurons via back propagation of action potentials (Fig. S3J-L).

## Discussion

In this study, using anatomical tracing techniques, we characterized in detail the whole-brain output pattern of aVMHvl^PR+^ neurons, showing projections to more than 150 brain regions, strongly projecting to hypothalamic, midbrain and thalamic regions. Moreover, our analysis revealed that the efferent connectivity of aVMHvl^PR+^ neurons presents shared and distinct output targets when compared to the efferent connectivity of pVMHvl^PR+^ neurons. One prominent difference was found in the PAG: whereas aVMHvl^PR+^ neurons sends strong projections across the whole AP axis of the dmPAG, dlPAG and lPAG, the axon terminals of pVMHvl^PR+^ are mostly present in the most posterior lPAG and the vlPAG regions. Using optogenetics, we found that the activation of aVMHvl^PR+^ neurons leads to the specific activation of dmPAG neurons and modulates the behavior of sexually receptive females during a sexual encounter by eliciting an increase in rejection behavior, partially recapitulating the behavioral phenotype observed when directly stimulation the somas of aVMHvl^PR+^ neurons.

The differences in connectivity of PR+ neurons across the AP axis of the VMHvl reported in the present study are largely consistent with those previously described for the Esr1-expressing population^28^. For instance, while posterior Esr1+ neurons engage more prominently in intra-hypothalamic connectivity, anterior Esr1+ neurons produce major additional outputs to the midbrain. Moreover, we have identified several areas that receive strong input from PR+ neurons, and that had been previously shown to receive no or few projections from the VMHvl^Esr+^ subpopulation, which may reflect specific output regions of the PR+ population. These input recipients include the ASO, the supraoptic nucleus (SO) and the peripeduncular nucleus (PP) for aVMHvl^PR+^ neurons; and the posterodorsal preoptic nucleus (PD) and Barrington’s nucleus (B) for pVMHvl^PR+^ neurons.These areas are involved in a wide array of different functions, ranging from the control of vasopressinergic systems (Riva 1999), to male and female sexual behavior^39–42^ and bladder control^43^. Notably, the PD has been shown to be selectively activated by ejaculation and lesions in this region reduce mounting behavior and delay ejaculation^44,45^, while the PP in female rats has been shown to play a functional role in the control of lordosis^41^. This highlights functionally meaningful output regions in the context of sexual behavior that are specific projections of the PR+ neurons.

These AP connectivity differences may underlie aspects of the functional differences that have been reported for the PR-expressing population throughout the VMHvl AP axis in the context of female sexual behavior^21,24^. Other spatially organized functional heterogeneities have been observed for other molecularly defined subpopulations of VMHvl neurons. For instance, while regulation of female sexual receptivity seems to be localized to the most posterior-lateral part of the VMHvl^19^, which neurons mainly express Cckar^23^, its posterior-medial division is involved in aggressive behavior toward intruders in mothers^19^. And while the pVMHvl^ER+,Npy2r–^ population was shown to be involved in sexual receptivity, the pVMHvl^ER+,Npy2r+^ population is involved in maternal aggression^22^.

In this study we focused on the projections from the aVMHvl^PR+^ neurons to the PAG. The PAG is a major midbrain hub for translating a vast array of anatomical inputs, including hypothalamic signals, into coordinated motor outputs underlying social behaviours, including defense and mating, through its projections to spinal premotor circuits^46,47^. The PAG is a very large and complex structure with widespread transcriptional and functional heterogeneity^48,49^. The dorsal, lateral and ventrolateral subregions of the PAG are known to control different behaviors ranging from escape and active avoidance from a threat^50–53^ (dorsomedial and dorsolateral PAG or dm/dlPAG), to mating, hunting and attack^54–59^ (lateral PAG or lPAG), and immobility, freezing, and lordosis^49,57,60–63^ (ventrolateral PAG or vlPAG). We observed differences in the output connectivity of PR+ neurons across the VMH AP axis, namely while the terminals of anterior neurons could be detected spanning the whole PAG (with prominent projections to the dmPAG, dlPAG and lPAG), posterior neurons terminate more posterior in lPAG and vlPAG. These differences in connectivity agree with the functional heterogeneity that has been reported for the PR+ population, as different subregions of the PAG are involved in the control of distinct behaviors.

The analysis of cFos density in the PAG after the optogenetic experiment revealed that artificial activation of aVMHvl^PR+^ neurons specifically activates the medial dmPAG. We observed a slight increase in cFos density in all PAG subdivisions, but it was more pronounced in the medial dmPAG, which is consistent with the previously proposed hypothesis that the PAG columns are co-activated during diverse behaviors^48^. In Vaughn et al.^64^, the authors investigate the transcriptional and spatial logic of PAG function during instinctive behaviors, not only across the different columns, but also along the entire AP axis. They observed that after mating, in the female brain, the more posterior part of PAG is active, while aggression in males (not studied in females) recruits the more anterior columns. This is consistent with our results that show that pVMHvl (involved in sexual receptivity) neurons project mainly to the posterior PAG, while the aVMHvl^PR+^ population projects strongly throughout the AP axis. These results suggest that the aVMHvl^PR+^-medial dmPAG pathway is a putative pathway for the control of sexual rejection behavior. We tested this hypothesis by optogenetically stimulating the axons of the aVMHvl^PR+^ neurons terminating in the medial dmPAG, in receptive females (in PE phase). Overall, the optogenetic manipulation resulted in disruptions of the sexual interaction, in particular, we observed an increase in the number of females that exhibited sexual rejections, similar to what we observed when we stimulated the somas of aVMHvl^PR+^ neurons^24^. However, contrary to the soma stimulation, we were not able to fully disrupt copulation, as all couples had sex and all the males ejaculated. While the soma stimulation leads to the activation of all projections from aVMHvl^PR+^, terminal activation was localized to the dmPAG. The fact that we did not observe any cFos activation in aVMHvl^PR+^ after the stimulation of its dmPAG terminals suggests that there was no backpropagation^65^, thus the manipulation was specific to the location where the optical fiber was present. However, definite confirmation would require pharmacologically silencing the silencing of the cell bodies while stimulating the axon terminals. As such, the most parsimonious explanation for the lesser behavioral disruption lies in the fact that in the present study we manipulated a single output of the aVMHvl^PR+^ population, the dmPAG. The more extreme phenotype after soma stimulation might have resulted from the fact that we did not manipulate other aVMHvl^PR+^ terminals in the PAG, or projections to other brain regions, such as other hypothalamic areas. Recent studies have shown that distinct projections of a neural population play separate roles in behavior. For instance, Liu et al., have shown that downstream pathways of the MPN mediate different aspects of social isolation^66^, and Lee et al., have shown that different DRN projections mediate distinct behaviors in social homeostasis, despite substantial collateralization^67^. Therefore, these results raise the question of whether there are other pathways involved in the regulation of rejection behavior or if it is specific to the aVMHvl^PR+^-dmPAG. We believe that complementary mechanisms and additional neural pathways are necessary for the full expression of the non-receptive state. In males, oxytocin neurons in the retrochiasmatic supraoptic nucleus (SO) were shown to be essential for social avoidance^68^. In addition, GABAergic neurons of the AHN have been identified to mediate anxiety-associated investigatory behaviors, during risk assessment, in the context of predatory threat^69^. In this study, we show that the aVMHvl^PR+^ neurons project to these nuclei, raising the hypothesis that the aVMHvl^PR+^-SO or aVMHvl^PR+^-AHN pathways could also be mediating social avoidance in females.

Altogether, our findings advance our understanding of the circuit logic through which aVMHvl^PR⁺^ neurons influence female sexual behavior. By linking the spatially organized outputs of these neurons to functionally distinct PAG subregions, this study identifies the aVMHvl^PR+^–dmPAG pathway as a candidate circuit component contributing to sexual rejection during non-receptive states within a larger, multi-node circuit.

## Method details

### Animals and and reproductive/estrous cycle monitoring

Data were collected from adult (2–9 months old) B6129S(Cg)-Pgrtm1.1(Cre)Shah/AndJ75 (PR-Cre^20^; JAX stock #017915), expressing Cre recombinase under the control of the PR promoter.

Animals were kept under controlled temperature of 23±1 °C, reversed photoperiod of 12 h light/dark cycle (light available from 8 pm to 8 am) and group-housed conditions (unless specified otherwise) in standard cages with environmental enrichment elements (cardboard igloo, shredded paper and soft nesting material). Food and water were provided *ad libitum*. Females were weaned at 20–21 days of age and group-housed with two to five animals. For optogenetic experiments, after reaching 6 weeks of age, females were exposed to adult C57BL/6 (JAX stock #000664) male soiled bedding once per week to stimulate the natural reproductive cycle. Cages were not changed on experimental days.

C57BL/6J male mice that had at least 3 prior ejaculation experiences within 3–4 weeks were used as studs for sexual behavior. For the training sessions, we used hormonally primed OVX females. The hormones were dissolved in sesame oil (Sigma-Aldrich, S3547-1L): estrogen (0,1 mg/ml, β-Estradiol 3-benzoate, Sigma-Aldrich, E8515-200MG) and progesterone (5 mg/ml, Sigma-Aldrich, P0130-25G). Females received 0,1 ml of estrogen 2 days prior to the sexual behavior and 0,1 ml of estrogen 4 h before the encounter, subcutaneously in the back while under anesthesia (with 3 % isoflurane in oxygen). All mice were kept in Specific Pathogen Free level 1 conditions.

Procedures were executed in accordance with the standards approved by the Commission for Experimentation and Animal Welfare of the Champalimaud Centre for the Unknown (Órgão para o Bem Estar Animal; ORBEA) and by the Portuguese National Authority for Animal Health (Direcção Geral de Alimentação e Veterinária; DGAV) (references 0421/000/000/2018 and 122571/24-S).

For the optogenetic experiments, females were habituated to Papanicolaou (Pap) smear collection daily for at least 4–5 days (approx. length of one complete reproductive cycle) before the experimental day. Pap smear collection consisted of manual restraint of the female followed by lavage of the vaginal region, using a pipette (Axygen, TE-204-Y, with a soft tip so that the vaginal region is not stimulated): 10 μL 0.1 mM PBS, which was then collected onto a slide^70^. These smears were then stained using the Papanicolaou staining protocol^70^, observed under a Zeiss AxioScope A1 brightfield microscope with a 10 x objective, and the reproductive state was determined by manual assessment of the proportion of different cell types in the stained smear^13,70–72^. Females were classified as D/diestrus or non-receptive when the predominant cell type was leukocytes, and as PE/between proestrus and estrus or receptive when there was a higher number of anucleated cornified cells along with an equal or lower proportion of nucleated epithelial cells in the smear. Experiments were performed immediately after pap smear collection and staining if the female was assessed as being in the PE or D state (specified in each experiment). Females that did not exhibit proper reproductive cycles during the habituation phase were not included in the study. All female mice used in this study were drug and test naive and had not been used for any previous procedures. Females had no previous sexual experience (sexually naive).

### Stereotaxic surgeries

For anatomical tracing, females were unilaterally injected on a stereotaxic (Kopf, Model 962) with 13.8 nL of AA1-CAG-Floxed-SynGFPrev-WPre (prepared in-house) in the aVMHvl (AP: −1.05, ML: –0.50, DV: −5.70; N = 5) or pVMHvl (AP: −1.3, ML: –0.70, DV: −6.00; N = 6) at 4.6 nL/pulse using a nanoliter injector (Nanoject II, Drummond Scientific, 3-000-205A) at 0.1 Hz. Females were perfused 3 weeks after viral injection.

For optogenetic activation of aVMHvl^PR+^ axons during male interactions, females were bilaterally injected with 100 nL of AAV9-EF1a-double floxed-hChR2(H134R)-EYFP-WPRE-HGHpA (Addgene, 20298) for the ChR2 group (N = 9) or 75 nL of AAV9-CAG-GFP (prepared in-house) for the Ctrl group (N = 7) in the aVMHvl (AP: −0.85, ML: ±0.50, DV: −5.60) at 4.6 nL/pulse using a nanoliter injector at 0.1 Hz. Subsequently, 400 μm diameter mono fiber-optic cannulas (Doric, MFC_400/430-0.48_3mm_SM3_FLT) were implanted above the medial dmPAG on a 26° angle (AP: −4.30, ML: +0.90, DV: −1.95) and were fixed using a light-cured dental cement (Optibond Universal, Kerr Dental, 36519) and a nano optimized fluid composite (Tetric Evoflow Universal, Ivoclar Vivadent, 595953).

A 300 μL intraperitoneal injection of saline and a 100 μL subcutaneous injection of buprenorphine was administered 0.5 h before the end of each surgery. Cages were left on a heating pad (Vet-Tech, HE008A) for 24 h. Females were thereafter singly housed and allowed to recover for at least 2 weeks before pap smear habituation was initiated.

### Electrophysiological recordings

For *ex vivo* electrophysiological recordings of PAG cells during optogenetic activation of aVMHvl^PR+^ axons, adult PR-Cre female mice (N = 4) were injected unilaterally with 150 nL of AAV9-EF1a-double floxed-hChR2(H134R)-EYFP-WPRE-HGHpA (Addgene, 20298) into the aVMHvl. Three weeks after viral injection, mice were deeply anesthetized using isofluorane and decapitated. Following decapitation, brains were quickly removed and placed in an “ice-cold” solution containing (in mM): 0.66 kynurenic acid, 3.63 pyruvate, 2.5 KCl, 1.25 NaH2PO4, 26 NaHCO3, 10 D-Glucose, 230 Sucrose, 0.5 CaCl2, 10 MgSO4, and bubbled with carbogen (5% CO2 and 95% O2). Coronal brain slices of 300 µm thickness containing the PAG were obtained using a vibratome (Leica, VT1200) and continuously perfused with oxygenated artificial cerebrospinal fluid (ACSF) containing (in mM): 127 NaCl, 2.5 KCl, 25 NaHCO3, 1.25 NaH2PO4, 25 D-Glucose, 2 CaCl2, and 1 MgCl2, at 34° C for 30 min and stored in the same solution at room temperature. Slices were then transferred to a recording chamber containing circulating oxygenated ACSF and visualized using a SliceScope Pro microscope (Scientifica). Whole-cell recordings were performed using pipettes (resistance 3-5 mV) filled with an internal solution containing (in mM): 135 K-gluconate, 10 HEPES, 10 Na-phosphocreatine, 3 Na-L-ascorbate, 4 MgCl2,4 Na-ATP, 0.4 Na-GTP (pH 7.2 adjusted with NaOH and osmolarity; 292 mOsm). Recordings were obtained with a Multiclamp 700B amplifier (Molecular Devices) and digitized at 10 KHz with a Digidata 1440a digitizer (Molecular Devices). Optogenetic activation was delivered using a 450 nm LED mounted on the back of the microscope (CoolLED, pR-300 Lite), with light being delivered to the slice through a 40 x objective lens at a focal distance of approximately 2 mm (Olympus, LUMPLFLN 40XW). Light pulses of 5 ms were delivered at 4 mW and 20 Hz.

### *In vivo* optogenetic manipulations

Light was delivered to the cannula using a 450 nm LED (Doric Lenses, LDFLS_450/075) via a 400 μm mono fiber-optic patch cords (Doric Lenses, D207-1405). Light stimulation (5 ms, 20 Hz, 3-4 mW) was delivered for 1 s every 5 s (1 s on – 4 s off) throughout the session, as used in other studies of optogenetic terminal stimulation of hypothalamic neurons^37,38^. Before each session, the LED power was measured at each tip of the patch cord using an optical power meter (Thorlabs, Inc., PM130D) and set to 3–4 mW.

All experiments were performed on PE females. The experimental session began with an habituation of 5 min in the experimental arena alone, followed by an open field test (OFT). Afterwards, a female was added into the arena and they were allowed to freely interact for 20 min. The female was removed and a male stud was added into the arena. The session was terminated either right after the male ejaculated, 30 min after the first mount attempt by the male if the female did not allow the male to successfully mount her until then, or 1 h after first mount with intromission if the male did not ejaculate until then. All our males were screened for sexual performance and we based the interval on previous studies of the lab^73,74^ where we observed that after the first mount attempt, most males ejaculated in less than 10 min. If the female did not allow the male to successfully mount her, a follow up pap smear was collected immediately after the conclusion of the experiment to assess if the female had transitioned to the next phase of the reproductive cycle, which would explain her non-receptive behavior. Only females that had follow-up pap smears still in the PE state were included. If the male did not exhibit a mount attempt within 30 min of introduction into the cage, he was removed and another male was added to the cage.

Video recording was performed using 2 cameras (Point Grey, Flea3, Monochrome), one top view and one front view, at 30 frames/second (fps). A custom data acquisition and synchronization board (constructed in-house) was used for triggering the LED and video cameras and was controlled using a custom program written in Bonsai 2.4.0.

On the day of euthanasia, females underwent 10 min of the experimental light stimulation protocol in the experimental cage before being returned to their home cage. Perfusion was performed after 90 min for cFos detection.

### Behavioral annotations

Behaviors during the sexual behavior assays were manually annotated frame-by-frame using Python Video Annotator 3.9.21 (Software Platform, Champalimaud Research). The following behavioral events were scored:

- Male entry - point event - when the male was added to the cage and all four of his paws hit the cage floor.
- Anogenital investigation - point event - when the nose of the animal is in the vicinity of the anogenital region of the other animal. This could be a male to female anogenital investigation event or vice-versa.
- Mount attempt - point event - when the male puts his paws on the back of the female in order to mount her, but fails to perform shallow thrusts (probing) or thrusting.
- Mount with intromission and intravaginal thrust - window of time - starts when the male puts his paws on the back of the female, same as a mount attempt, and is able to perform intromission and intravaginal thrusts. Ends when the male removes his paws from the back of the female.
- Ejaculation - point event - when the male enters into the last thrust before shivering and falling to the side.
- Rejection - point event - when the female lifts her hind- or fore-paw to hit the male.
- Escape - point event - when the female begins to run away from a mount attempt of a male and is successful in doing so within 15 frames of the mount attempt.

The distance between the two animals was calculated using the frame-by-frame 2-body centroid tracking node of Bonsai.

### Histology

Females were deeply anesthetized with sodium pentobarbital (120 mg/kg of body weight) and perfused transcardially using 0.1 M PBS followed by fixation using ice-cold 0.4% paraformaldehyde in 0.1 M PBS. The brains were removed and stored in 30% sucrose-PBS + 0.1% sodium azide solution until being sliced. The heads of the females used in the optogenetic experiments were severed and stored whole at 4℃ in 0.4% PFA for 24–48 h before the brains were removed. Brains were coronally sliced on a sliding microtome (Leica, SM2000 R) or a cryostat (Leica, CM1800) at 45–50 μm thickness.

Anatomical tracing: The brain sections required no immunostaining, as the SynGFP exhibits strong native fluorescence. After sliced, the brain sections were mounted on glass slides (Thermo Fisher Scientific) using Mowiol (Sigma-Aldrich) as a mounting medium, cover-slipped (Thermo Scientific Menzel) and kept at 4℃ until imaging.

Optogenetics experiment: floating brain slices underwent a double immunostaining for EYFP and cFos using a primary antibody cocktail of rabbit anti-cFos (1:2000, Synaptic Systems, Cat. No. 226 008) and goat anti-GFP (1:1000, Abcam, ab6673) followed by a secondary antibody cocktail of Alexa Fluor 594 donkey anti-rabbit (1:1000, Invitrogen, ab150076) and Alexa Fluor Plus 488 donkey anti-goat (1:1000, Invitrogen, A32814).

Slices were mounted on glass slides (Thermo Fisher Scientific) using Mowiol (Sigma-Aldrich) as a mounting medium, cover-slipped (Thermo Scientific Menzel) and kept at 4℃ until imaging.

### Imaging

Anatomical tracing: The whole-brain sections were imaged using a slide scanner (Zeiss Microscopy, ZeissAxioScan.Z1).

Optogenetics experiment. For brain sections in which colocalization analysis was conducted, a confocal laser scanning microscope with Airyscan (Zeiss, LSM 710) equipped with a 10 x/0.45 apochromat lens (Zeiss) was used for acquisition. Brain slices containing the aVMHvl were identified by overlaying the Allen brain atlas^75^ (© 2004 Allen Institute for Brain Science. Allen Mouse Brain Atlas) on brain slice images in Adobe Illustrator 2020 (Adobe).

### Quantification and statistical analysis

#### Brain section image analysis of Synaptophysin-GFP

The obtained images were processed using the Zen software (Zen 2.6, Zeiss Microscopy). Shading correction and stitching were applied and finally images were resized to 50% and exported as portable network graphics (.png).

The images were then analyzed following the QUINT workflow^76^ with some adaptations. The process involved the following steps:

1. Pre-processing: Images were processed in Nutil using the Resize function to meet the QuickNII input requirement of a maximum image size of 16 MP. The same resize factor was applied to all images within a batch, ensuring all images met the requirements.
2. Atlas registration: For each animal, an extensible markup language (XML) descriptor file containing all its brain sections was generated using the “FileBuider.bat” tool provided with QuickNII (RRID:SCR_016865). Image registration to the Allen brain atlas as reference for the brain regions (© 2004 Allen Institute for Brain Science. Allen Mouse Brain Atlas)^75^ was performed using QuickNII.
3. Segmentation: Segmentation of labelled features was performed using ilastik. The segmentation was performed in two steps. First, using Pixel Classification, as described in Yates et al. (2019)^76^, obtaining a probability map. The second step would be Object Classification, however it was not appropriate for our type of signal (puncta). To overcome this, we used a customized MATLAB script to save the segmentation images obtained in the previous step. A probability threshold of 0.55 was applied to first binarize the image, and then the image was saved as RGB color mode, with the signal in the red channel only.
4. Quantification: Quantification of features per atlas region was performed by using the Quantifier feature of Nutil, as described^76^. This step generated a set of report files and customized atlas images superimposed with the object pixels (signal).

Customized MATLAB scripts were used to analyze the data of the reports. Several exclusion criteria were applied in sequential order: (1) areas that had 0 object pixels for all animals of the batch were excluded; (2) brain regions that 4 out of 5 animals (for aVMHvl) or 5 out of 6 animals (for pVMHvl) had 0 object pixels were excluded; (3) brain regions that had less than 9 object pixels in more than 4 animals (for both batches), it was considered below signal and those areas were excluded. After this, brain areas with subregions were clustered together, summing the object pixels and the area (e.g. ‘Primary motor area, Layer 2/3’, ‘Primary motor area, Layer 5’, ‘Primary motor area, Layer 6a’, ‘Primary motor area, Layer 6b’ were clustered to ‘Primary motor area’). (4) Signal on fiber tracts was not included in the analysis.

As a result, we obtained the area (in pixels) and the signal (in pixels) for each brain region. By determining the pixel size in μm2, per batch, we calculated the area in μm2 and the density of the signal. Finally, the density was normalized to the number of somas in the injection site ((# of fluorescent pixels/area (mm2))/# of somas). The number of somas was manually counted, using the images with their original size.

For the detailed analysis of Synaptophysin-GFP in the periaqueductal gray (PAG), brain sections spanning the full AP extent of the PAG were selected. The number of AP levels was defined by the number of slices of the animal that contained less sections (per batch). Slices were selected so they would match between all animals (per AP level). If there were missing slices, the synGFP density values were interpolated and plotted as a function of AP levels, in MATLAB. Bregma value per section was obtained from QuickNII.

To generate the PAG heatmaps, we performed image registration in MATLAB to align the PAG slices. We used the segmentation images obtained in step 3 of the adapted QUINT workflow, as previously described. Per anteroposterior level referenced in Bregma, the segmentation map of one animal was fixed and used as reference. To the segmentation maps from the other animals a geometric transformation to the reference image was applied. With that, we obtained heatmaps with the SynGFP signal (in pixels) for each AP level that had the same geometric position. The signal was binned, normalized to the number of somas in the injection site, averaged between animals and plotted as an image.

### Behavioral analysis for optogenetic experiments

We tracked the position of the animals’ center of mass using Bonsai. The distance between the couple, duration of the phases and rate of events were calculated using Python. Box plots were generated using the *boxplot* function, scatter plots were generated using the *plot* function and the survival analysis was calculated using the *KaplanMeierFitter* in Python.

### cFos quantification

The location of brain regions was manually determined by overlaying vectorized Allen Brain Atlas (© 2004 Allen Institute for Brain Science. Allen Mouse Brain Atlas)^75^ reference sections in Adobe Illustrator (Adobe).

cFos positive cells were manually counted in aVMHvl and PAG and normalized to the area, calculated in ImageJ 1.52n. Two brain slices containing aVMHvl from each animal were used for the quantification of cFos expression. Three brain slices containing PAG (one anterior (∼ -3.40 to -3.52 mm), one medial (∼ -4.36 to -4.60 mm) and one posterior (∼ -4.96 to -5.02 mm)) from each animal were used for the quantification cFos expression, using Paxino’s Mouse Brain Atlas as reference^77^. Cell counting was performed manually using ImageJ.

### Statistical analysis

Statistical analyses were performed using GraphPad Prism 9 Software. Normality of the residuals was tested using the Shapiro-Wilk test. In data that passed the normality test, independent-samples Student’s t-test or paired t-test were used to evaluate differences between groups. If data did not follow a normal distribution, analysis was performed using a non-parametric Mann–Whitney U test for unpaired samples.

Box plots indicate median and the interquartile range (IQR, ± 25th – 75th percentile) and the whisker edges represent the minimum and maximum data limits excluding outliers using the Tukey criterion (outliers are depicted outside the box plot). Error bars and shaded error bars represent mean and standard error of the mean. For comparing fractions (i.e. percentage of animals that had sex or rejected), we used the N-1 Two-proportion test. For the survival function analysis, we used the Cox proportional-hazards model. Statistical significance was set at p < 0.05. Statistic details (test and p values) can be found in the results section.

All mice were randomly assigned into experimental groups. Each experimental batch consisted of age and weight matched groups of females. Each experimental batch was balanced between control and test groups. Blinding of the data was conducted before any manual scoring (both for behavioral annotations and for cFos quantification). No sample size estimation was conducted prior to this study.

For optogenetics experiments, biological replicates consisted of independent mice (N = number of mice) tested only once. These results were analyzed with unpaired statistics. For neural recording and manipulation experiments, mice were excluded only when virus or fiber location was visibly off-target.

## References

1. Clutton-brock, T.H., and Parker, G.A. (1995). Sexual coercion in animal societies. Anim. Behav. 49, 1345–1365. 10.1006/anbe.1995.0166.

2. Clutton-Brock, T.H., and Parker, G.A. (1995). Punishment in animal societies. Nature 373, 209–216. 10.1038/373209a0.

3. Johnstone, R.A., and Keller, L. (2000). How Males Can Gain by Harming Their Mates: Sexual Conflict, Seminal Toxins, and the Cost of Mating. Am. Nat. 156, 368–377. 10.1086/303392.

4. Constantz, G.D., and Smith, R.L. (1984). Sperm Competition and the Evolution of Animal Mating Systems (Elsevier) 10.1016/B978-0-12-652570-0.X5001-5.

5. Parker, G.A. (2006). Sexual conflict over mating and fertilization: an overview. Philos. Trans. R. Soc. B Biol. Sci. 361, 235–259. 10.1098/rstb.2005.1785.

6. Garratt, M., Try, H., Smiley, K.O., Grattan, D.R., and Brooks, R.C. (2020). Mating in the absence of fertilization promotes a growth-reproduction versus lifespan trade-off in female mice. Proc. Natl. Acad. Sci. 117, 15748–15754. 10.1073/pnas.2003159117.

7. Krenhardt, K., Markó, G., Jablonszky, M., Török, J., and Garamszegi, L.Z. (2021). Sex-dependent risk-taking behaviour towards different predatory stimuli in the collared flycatcher. Behav. Processes 186, 104360. 10.1016/j.beproc.2021.104360.

8. Tybur, J.M., and Gangestad, S.W. (2011). Mate preferences and infectious disease: theoretical considerations and evidence in humans. Philos. Trans. R. Soc. B Biol. Sci. 366, 3375–3388. 10.1098/rstb.2011.0136.

9. Yin, L., and Lin, D. (2023). Neural control of female sexual behaviors. Horm. Behav. 151. 10.1016/j.yhbeh.2023.105339.

10. Dias, I.C., Gutierrez-Castellanos, N., Lenschow, C., and Lima, S.Q. (2025). Ready or not: Neural mechanisms regulating female sexual behavior. Curr. Opin. Neurobiol. 93, 103069. 10.1016/j.conb.2025.103069.

11. Sutton Hickey, A.K., and Krashes, M.J. (2020). Integrating Hunger with Rival Motivations. Trends Endocrinol. Metab. TEM 31, 495–507. 10.1016/j.tem.2020.04.006.

12. Geraghty, A.C., Muroy, S.E., Zhao, S., Bentley, G.E., Kriegsfeld, L.J., and Kaufer, D. (2015). Knockdown of hypothalamic RFRP3 prevents chronic stress-induced infertility and embryo resorption. eLife 4, e04316. 10.7554/eLife.04316.

13. Gutierrez-Castellanos, N., Husain, B.F.A., Dias, I.C., and Lima, S.Q. (2022). Neural and behavioral plasticity across the female reproductive cycle. Trends Endocrinol. Metab. 33, 769–785. 10.1016/j.tem.2022.09.001.

14. Pfaff, D.W., and Sakuma, Y. (1979). Facilitation of the lordosis reflex of female rats from the ventromedial nucleus of the hypothalamus. J. Physiol. 288, 189–202. 10.1113/jphysiol.1979.sp012690.

15. Pfaff, D.W., and Sakuma, Y. (1979). Deficit in the lordosis reflex of female rats caused by lesions in the ventromedial nucleus of the hypothalamus. J. Physiol. 288, 203–210. 10.1113/jphysiol.1979.sp012691.

16. Rubin, B.S., and Barfield, R.J. (1980). Priming of Estrous Responsiveness by Implants of 17 *β* - Estradiol in the Ventromedial Hypothalamic Nucleus of Female Rats*. Endocrinology 106, 504–509. 10.1210/endo-106-2-504.

17. Rubin, B.S., and Barfield, R.J. (1983). Progesterone in the Ventromedial Hypothalamus Facilitates Estrous Behavior in Ovariectomized, Estrogen-Primed Rats*. Endocrinology 113, 797–804. 10.1210/endo-113-2-797.

18. Rissman, E.F., Early, A.H., Taylor, J.A., Korach, K.S., and Lubahn, D.B. (1997). Estrogen receptors are essential for female sexual receptivity. Endocrinology 138, 507–510. 10.1210/endo.138.1.4985.

19. Hashikawa, K., Hashikawa, Y., Tremblay, R., Zhang, J., Feng, J.E., Sabol, A., Piper, W.T., Lee, H., Rudy, B., and Lin, D. (2017). Esr1+ cells in the ventromedial hypothalamus control female aggression. Nat. Neurosci. 20, 1580–1590. 10.1038/nn.4644.

20. Yang, C.F., Chiang, M.C., Gray, D.C., Prabhakaran, M., Alvarado, M., Juntti, S.A., Unger, E.K., Wells, J.A., and Shah, N.M. (2013). Sexually Dimorphic Neurons in the Ventromedial Hypothalamus Govern Mating in Both Sexes and Aggression in Males. Cell 153, 896–909. 10.1016/j.cell.2013.04.017.

21. Inoue, S., Yang, R., Tantry, A., Davis, C., Yang, T., Knoedler, J.R., Wei, Y., Adams, E.L., Thombare, S., Golf, S.R., et al. (2019). Periodic Remodeling in a Neural Circuit Governs Timing of Female Sexual Behavior. Cell 179, 1393–1408.e16. 10.1016/j.cell.2019.10.025.

22. Liu, M., Kim, D.-W., Zeng, H., and Anderson, D.J. (2022). Make war not love: The neural substrate underlying a state-dependent switch in female social behavior. Neuron 110, 841–856.e6. 10.1016/j.neuron.2021.12.002.

23. Yin, L., Hashikawa, K., Hashikawa, Y., Osakada, T., Lischinsky, J.E., Diaz, V., and Lin, D. (2022). VMHvllCckar cells dynamically control female sexual behaviors over the reproductive cycle. Neuron 110, 3000–3017.e8. 10.1016/j.neuron.2022.06.026.

24. Gutierrez-Castellanos, N., Husain, B.F.A., Dias, I.C., Nomoto, K., Duarte, M.A., Ferreira, L., Lacoste, B., and Lima, S.Q. (2024). A hypothalamic node for the cyclical control of female sexual rejection. Neuron. 10.1016/j.neuron.2024.10.026.

25. Swanson, L.W. (2000). Cerebral hemisphere regulation of motivated behavior. Brain Res. 886, 113–164. 10.1016/s0006-8993(00)02905-x.

26. Swanson, L.W. (2005). Anatomy of the soul as reflected in the cerebral hemispheres: neural circuits underlying voluntary control of basic motivated behaviors. J. Comp. Neurol. 493, 122–131. 10.1002/cne.20733.

27. Luiten, P.G., ter Horst, G.J., and Steffens, A.B. (1987). The hypothalamus, intrinsic connections and outflow pathways to the endocrine system in relation to the control of feeding and metabolism. Prog. Neurobiol. 28, 1–54. 10.1016/0301-0082(87)90004-9.

28. Lo, L., Yao, S., Kim, D.-W., Cetin, A., Harris, J., Zeng, H., Anderson, D.J., and Weissbourd, B. (2019). Connectional architecture of a mouse hypothalamic circuit node controlling social behavior. Proc. Natl. Acad. Sci. 116, 7503–7512. 10.1073/pnas.1817503116.

29. Blaustein, J.D., and Turcotte, J.C. (1989). Estradiol-induced progestin receptor immunoreactivity is found only in estrogen receptor-immunoreactive cells in guinea pig brain. Neuroendocrinology 49, 454–461. 10.1159/000125152.

30. Sá, S.I., and Fonseca, B.M. (2017). Dynamics of progesterone and estrogen receptor alpha in the ventromedial hypothalamus. J. Endocrinol. 233, 197–207. 10.1530/JOE-16-0663.

31. Sürmeli, G., Marcu, D.C., McClure, C., Garden, D.L.F., Pastoll, H., and Nolan, M.F. (2015). Molecularly Defined Circuitry Reveals Input-Output Segregation in Deep Layers of the Medial Entorhinal Cortex. Neuron 88, 1040–1053. 10.1016/j.neuron.2015.10.041.

32. Estela-Pro, V.J., and Burwell, R.D. (2022). The anatomy and function of the postrhinal cortex. Behav. Neurosci. 136, 101–113. 10.1037/bne0000500.

33. Massey, P.V., and Bashir, Z.I. (2007). Long-term depression: multiple forms and implications for brain function. Trends Neurosci. 30, 176–184. 10.1016/j.tins.2007.02.005.

34. Sheridan, N., and Tadi, P. (2025). Neuroanatomy, Thalamic Nuclei. In StatPearls (StatPearls Publishing).

35. Vertes, R.P., Linley, S.B., Groenewegen, H.J., and Witter, M.P. (2015). Chapter 16 - Thalamus. In The Rat Nervous System (Fourth Edition), G. Paxinos, ed. (Academic Press), pp. 335–390. 10.1016/B978-0-12-374245-2.00016-4.

36. Canteras, N.S., Simerly, R.B., and Swanson, L.W. (1994). Organization of projections from the ventromedial nucleus of the hypothalamus: A *Phaseolus vulgaris* -Leucoagglutinin study in the rat. J. Comp. Neurol. 348, 41–79. 10.1002/cne.903480103.

37. Ahmadlou, M., Giannouli, M., Van Vierbergen, J.F.M., Van Leeuwen, T., Bloem, W., Houba, J.H.W., Shirazi, M.Y., Cazemier, J.L., Haak, R., Dubey, M., et al. (2024). Cell-type-specific hypothalamic pathways to brainstem drive context-dependent strategies in response to stressors. Curr. Biol. 34, 2448–2459.e4. 10.1016/j.cub.2024.04.053.

38. Cao, M., Ammari, R., Chen, M.X., Wai, P., Sahni, A., Liang, S., Legrave, N., Macrae, J., Strom, M., and Kohl, J. (2024). Integration of hunger and hormonal state gates infant-directed aggression. Preprint at bioRxiv, 10.1101/2024.11.25.625278 https://doi.org/10.1101/2024.11.25.625278.

39. Simmons, D.A., and Yahr, P. (2002). Projections of the posterodorsal preoptic nucleus and the lateral part of the posterodorsal medial amygdala in male gerbils, with emphasis on cells activated with ejaculation. J. Comp. Neurol. 444, 75–94. 10.1002/cne.10128.

40. Hull, E.M., and Rodríguez-Manzo, G. (2009). Male sexual behavior. In Hormones, brain and behavior, Vol. 1, 2nd ed (Elsevier Academic Press), pp. 5–65. 10.1016/B978-008088783-8.00001-2.

41. López, H.S., and Carrer, H.F. (1985). Evidence for peripeduncular neurons having a role in the control of feminine sexual behavior in the rat. Physiol. Behav. 35, 205–208. 10.1016/0031-9384(85)90337-3.

42. Hansen, S., and Köhler, C. (1984). The importance of the peripeduncular nucleus in the neuroendocrine control of sexual behavior and milk ejection in the rat. Neuroendocrinology 39, 563–572. 10.1159/000124038.

43. Verstegen, A.M.J., Klymko, N., Zhu, L., Mathai, J.C., Kobayashi, R., Venner, A., Ross, R.A., VanderHorst, V.G., Arrigoni, E., Geerling, J.C., et al. (2019). Non-Crh Glutamatergic Neurons in Barrington’s Nucleus Control Micturition via Glutamatergic Afferents from the Midbrain and Hypothalamus. Curr. Biol. CB 29, 2775–2789.e7. 10.1016/j.cub.2019.07.009.

44. Heeb, M.M., and Yahr, P. (1996). C-Fos immunoreactivity in the sexually dimorphic area of the hypothalamus and related brain regions of male gerbils after exposure to sex-related stimuli or performance of specific sexual behaviors. Neuroscience 72, 1049–1071. 10.1016/0306-4522(95)00602-8.

45. Heeb, M.M., and Yahr, P. (2000). Cell-body lesions of the posterodorsal preoptic nucleus or posterodorsal medial amygdala, but not the parvicellular subparafascicular thalamus, disrupt mating in male gerbils. Physiol. Behav. 68, 317–331. 10.1016/S0031-9384(99)00182-1.

46. Stempel, A.V. (2024). A conserved brainstem region for instinctive behaviour control: The vertebrate periaqueductal gray. Curr. Opin. Neurobiol. 86, 102878. 10.1016/j.conb.2024.102878.

47. Zhang, H., Zhu, Z., Ma, W.-X., Kong, L.-X., Yuan, P.-C., Bu, L.-F., Han, J., Huang, Z.-L., and Wang, Y.-Q. (2024). The contribution of periaqueductal gray in the regulation of physiological and pathological behaviors. Front. Neurosci. 18, 1380171. 10.3389/fnins.2024.1380171.

48. Vianna, D.M.L., and Brandão, M.L. (2003). Anatomical connections of the periaqueductal gray: Specific neural substrates for different kinds of fear. Braz. J. Med. Biol. Res. 36, 557–566. 10.1590/S0100-879X2003000500002.

49. Vianna, D.M.L., Landeira-Fernandez, J., and Brandão, M.L. (2001). Dorsolateral and ventral regions of the periaqueductal gray matter are involved in distinct types of fear. Neurosci. Biobehav. Rev. 25, 711–719. 10.1016/S0149-7634(01)00052-5.

50. Evans, D.A., Stempel, A.V., Vale, R., Ruehle, S., Lefler, Y., and Branco, T. (2018). A synaptic threshold mechanism for computing escape decisions. Nature 558, 590–594. 10.1038/s41586-018-0244-6.

51. Lefler, Y., Campagner, D., and Branco, T. (2020). The role of the periaqueductal gray in escape behavior. Curr. Opin. Neurobiol. 60, 115–121. 10.1016/j.conb.2019.11.014.

52. Masferrer, M., Silva, B.A., Nomoto, K., Lima, S.Q., and Gross, C.T. (2020). Differential Encoding of Predator Fear in the Ventromedial Hypothalamus and Periaqueductal Grey. J. Neurosci. 40, 9283–9292. 10.1523/JNEUROSCI.0761-18.2020.

53. Stempel, A.V., Evans, D.A., Arocas, O.P., Claudi, F., Lenzi, S.C., Kutsarova, E., Margrie, T.W., and Branco, T. (2024). Tonically active GABAergic neurons in the dorsal periaqueductal gray control instinctive escape in mice. Curr. Biol. 34, 3031–3039.e7. 10.1016/j.cub.2024.05.068.

54. Falkner, A.L., Wei, D., Song, A., Watsek, L.W., Chen, I., Chen, P., Feng, J.E., and Lin, D. (2020). Hierarchical Representations of Aggression in a Hypothalamic-Midbrain Circuit. Neuron 106, 637–648.e6. 10.1016/j.neuron.2020.02.014.

55. Han, W., Tellez, L.A., Rangel, M.J., Motta, S.C., Zhang, X., Perez, I.O., Canteras, N.S., Shammah-Lagnado, S.J., Pol, A.N. van den, and Araujo, I.E. de (2017). Integrated Control of Predatory Hunting by the Central Nucleus of the Amygdala. Cell 168, 311–324.e18. 10.1016/j.cell.2016.12.027.

56. Rossier, D., La Franca, V., Salemi, T., Natale, S., and Gross, C.T. (2021). A neural circuit for competing approach and defense underlying prey capture. Proc. Natl. Acad. Sci. U. S. A. 118, e2013411118. 10.1073/pnas.2013411118.

57. Tovote, P., Esposito, M.S., Botta, P., Chaudun, F., Fadok, J.P., Markovic, M., Wolff, S.B.E., Ramakrishnan, C., Fenno, L., Deisseroth, K., et al. (2016). Midbrain circuits for defensive behaviour. Nature 534, 206–212. 10.1038/nature17996.

58. Yamada, S., and Kawata, M. (2014). Identification of neural cells activated by mating stimulus in the periaqueductal gray in female rats. Front. Neurosci. 8. 10.3389/fnins.2014.00421.

59. Yu, H., Xiang, X., Chen, Z., Wang, X., Dai, J., Wang, X., Huang, P., Zhao, Z.-D., Shen, W.L., and Li, H. (2021). Periaqueductal gray neurons encode the sequential motor program in hunting behavior of mice. Nat. Commun. 12, 6523. 10.1038/s41467-021-26852-1.

60. Lonstein, J.S., and Stern, J.M. (1998). Site and behavioral specificity of periaqueductal gray lesions on postpartum sexual, maternal, and aggressive behaviors in rats. Brain Res. 804, 21–35. 10.1016/S0006-8993(98)00642-8.

61. Sakuma, Y., and Pfaff, D.W. (1979). Facilitation of female reproductive behavior from mesensephalic central gray in the rat. Am. J. Physiol.-Regul. Integr. Comp. Physiol. 10.1152/ajpregu.1979.237.5.R278.

62. Sakuma, Y., and Pfaff, D.W. (1980). Covergent effects of lordosis-relevant somatosensory and hypothalamic influences on central gray cells in the rat mesencephalon. Exp. Neurol. 70, 269–281. 10.1016/0014-4886(80)90026-6.

63. St. Laurent, R., Martinez Damonte, V., Tsuda, A.C., and Kauer, J.A. (2020). Periaqueductal Gray and Rostromedial Tegmental Inhibitory Afferents to VTA Have Distinct Synaptic Plasticity and Opiate Sensitivity. Neuron 106, 624–636.e4. 10.1016/j.neuron.2020.02.029.

64. Vaughn, E., Eichhorn, S., Jung, W., Zhuang, X., and Dulac, C. (2022). Three-dimensional Interrogation of Cell Types and Instinctive Behavior in the Periaqueductal Gray. Preprint at bioRxiv, 10.1101/2022.06.27.497769 https://doi.org/10.1101/2022.06.27.497769.

65. Allen, B.D., Singer, A.C., and Boyden, E.S. (2015). Principles of designing interpretable optogenetic behavior experiments. Learn. Mem. 22, 232–238. 10.1101/lm.038026.114.

66. Liu, D., Rahman, M., Johnson, A., Amo, R., Tsutsui-Kimura, I., Sullivan, Z.A., Pena, N., Talay, M., Logeman, B.L., Finkbeiner, S., et al. (2025). A hypothalamic circuit underlying the dynamic control of social homeostasis. Nature 640, 1000–1010. 10.1038/s41586-025-08617-8.

67. Lee, C.R., Matthews, G.A., Lemieux, M.E., Wasserlein, E.M., Borio, M., Miranda, R.L., Keyes, L.R., Schneider, G.P., Jia, C., Tran, A., et al. (2025). Separable Dorsal Raphe Dopamine Projections Mimic the Facets of a Loneliness-like State. eLife. 10.7554/eLife.105955.2.

68. Osakada, T., Yan, R., Jiang, Y., Wei, D., Tabuchi, R., Dai, B., Wang, X., Zhao, G., Wang, C.X., Liu, J.-J., et al. (2024). A dedicated hypothalamic oxytocin circuit controls aversive social learning. Nature 626, 347–356. 10.1038/s41586-023-06958-w.

69. Hong, C.Y., Din, J.S., Chang, H., Bang, J.Y., and Kim, J.C. (2025). Anterior hypothalamic nucleus drives distinct defensive responses through cell-type-specific activity. iScience 28, 112097. 10.1016/j.isci.2025.112097.

70. Caligioni, C.S. (2009). Assessing Reproductive Status/Stages in Mice. Curr. Protoc. Neurosci. 48. 10.1002/0471142301.nsa04is48.

71. Pfaus, J.G. (1999). Neurobiology of sexual behavior. Curr. Opin. Neurobiol. 9, 751–758. 10.1016/S0959-4388(99)00034-3.

72. Snoeren, E.M.S. (2018). Female Reproductive Behavior. In Neuroendocrine Regulation of Behavior Current Topics in Behavioral Neurosciences., L. M. Coolen and D. R. Grattan, eds. (Springer International Publishing), pp. 1–44. 10.1007/7854_2018_68.

73. Nomoto, K., Ikumi, M., Otsuka, M., Asaba, A., Kato, M., Koshida, N., Mogi, K., and Kikusui, T. (2018). Female mice exhibit both sexual and social partner preferences for vocalizing males. Integr. Zool. 13, 735–744. 10.1111/1749-4877.12357.

74. Valente, S., Marques, T., and Lima, S.Q. (2021). No evidence for prolactin’s involvement in the post-ejaculatory refractory period. Commun. Biol. 4, 10. 10.1038/s42003-020-01570-4.

75. Lein, E.S., Hawrylycz, M.J., Ao, N., Ayres, M., Bensinger, A., Bernard, A., Boe, A.F., Boguski, M.S., Brockway, K.S., Byrnes, E.J., et al. (2007). Genome-wide atlas of gene expression in the adult mouse brain. Nature 445, 168–176. 10.1038/nature05453.

76. Yates, S.C., Groeneboom, N.E., Coello, C., Lichtenthaler, S.F., Kuhn, P.-H., Demuth, H.-U., Hartlage-Rübsamen, M., Roßner, S., Leergaard, T., Kreshuk, A., et al. (2019). QUINT: Workflow for Quantification and Spatial Analysis of Features in Histological Images From Rodent Brain. Front. Neuroinformatics 13, 75. 10.3389/fninf.2019.00075.

77. Franklin, K., and Paxinos, G. (2008). The Mouse Brain in Stereotaxic Coordinates, Compact (Elsevier Science).

