## Supplementary material for "Neural substrates of female sexual rejection: hypothalamic pathways to the periaqueductal gray": Supp data

### Supplementary Data

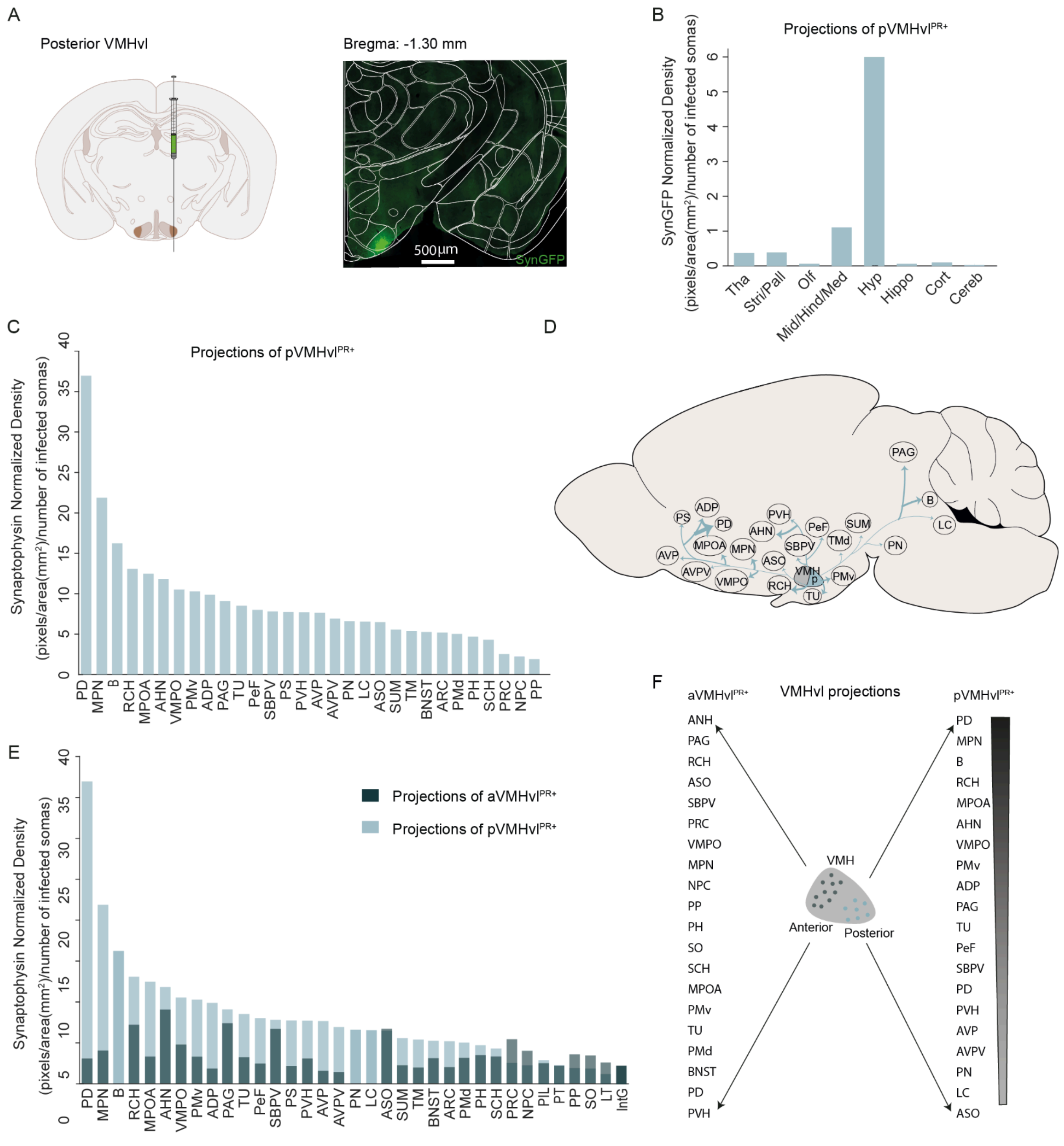

**Supplementary figure 1**, **related to Figure 1 - Outputs of posterior VMHvl$^{PR+}$ neurons.** (A) Schematic of pVMHvl virus injection (left) and representative histology image showing PR-Cre neurons expressing Synaptophysin-GFP (SynGFP) at the injection site (right). Scale bar: 500 μm. (B) Broad quantification of pVMHvl$^{PR+}$ projections to the main brain divisions: thalamus (Tha), striatum/pallidus (Stri/Pall), olfactory (Olf), midbrain/hindbrain/medulla (Mid/Hind/Med), hypothalamus (Hyp), hippocampus (Hippo), cortex (Cort), cerebellum (Cereb). (C) Quantification of the projections of pVMHvl$^{PR+}$ neurons. The analysis of the signal across 30 principal targets revealed strong projections from the pVMHvlPR+ neurons project strongly within the hypothalamus, including several areas of the periventricular regions, such as the posterodorsal preoptic nucleus (PD), ventromedial preoptic nucleus (VMPO), anterodorsal preoptic nucleus (ADP), subparaventricular zone (SBPV), parastrial nucleus (PS), anteroventral preoptic nucleus (AVP) and anteroventral periventricular nucleus (AVPV), and others including medial preoptic nucleus (MPN), retrochiasmatic area (RCH), medial preoptic area (MPOA), anterior hypothalamic nucleus (AHN), ventral premammillary nucleus (PMv); the pVMHvl$^{PR+}$ neurons also strongly project to the hindbrain regions, such as the Barrington's nucleus (B) and locus coeruleus (LC); as well as midbrain regions, including the periaqueductal grey (PAG) and the paranigral nucleus (PN). (D) Summary schematic with the main output regions of the pVMHvl$^{PR+}$ neurons. Arrow thickness represents relative projection strength. (E) Overlap of the quantification of the projections of both subpopulations for comparison. (F) Summary of the twenty main output regions of the aVMHvl$^{PR+}$ and pVMHvl$^{PR+}$ neurons in a descending order (top to bottom). Dark cyan: aVMHvl$^{PR+}$ neurons, N = 5; Light cyan: pVMHvl$^{PR+}$ neurons, N = 6.

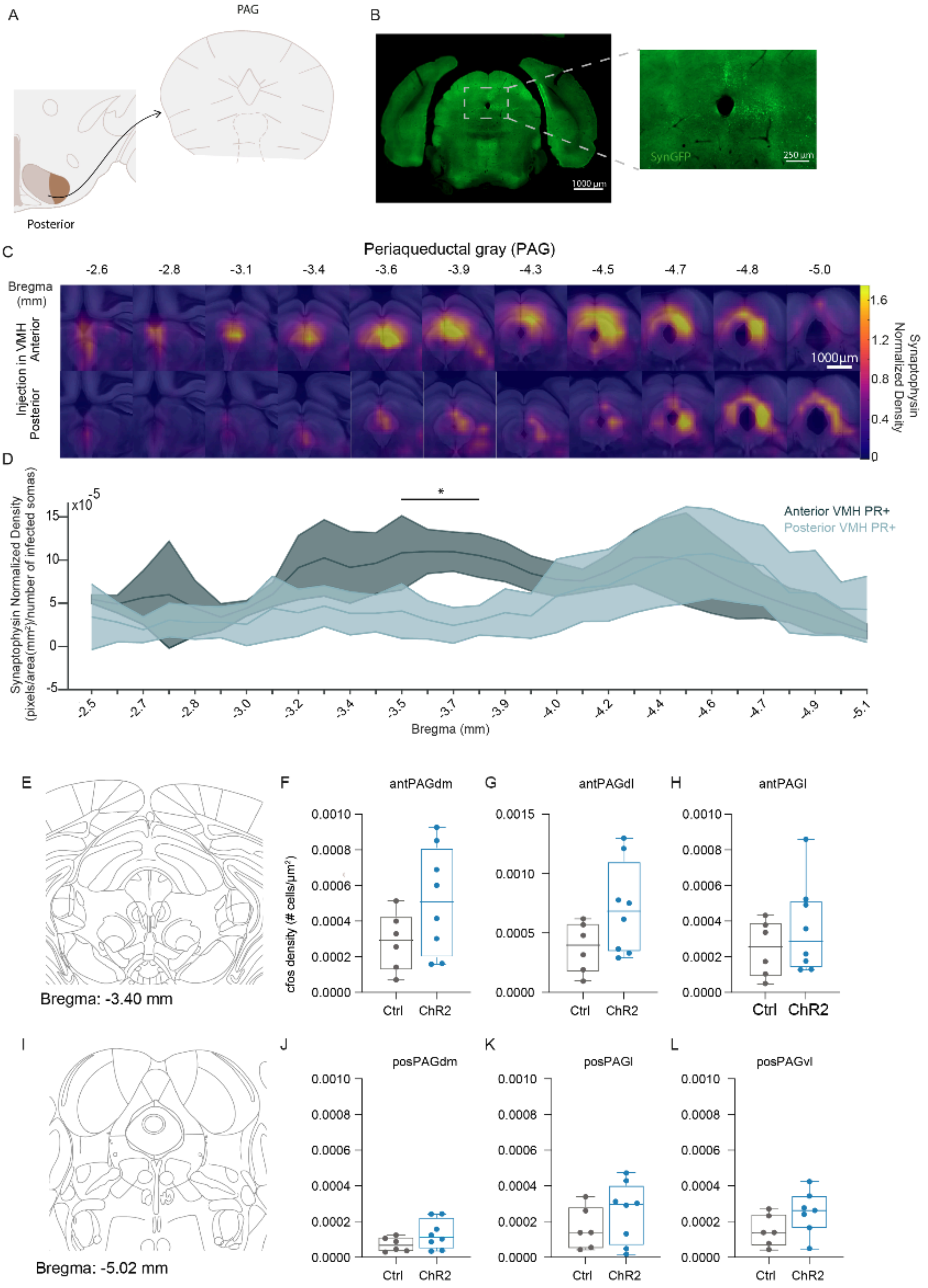
A
PAG
Posterior
B
SynGFP
250 µm
1000 µm
C
Periaqueductal gray (PAG)
Bregma (mm)
-2.6
-2.8
-3.1
-3.4
-3.6
-3.9
-4.3
-4.5
-4.7
-4.8
-5.0
Injection in VMH
Anterior
Posterior
1000µm
Synaptophysin Normalized Density
1.6
1.2
0.8
0.4
0
D
Synaptophysin Normalized Density (pixels/area(mm²)/number of infected somas)
x10-5
*
Anterior VMH PR+
Posterior VMH PR+
Bregma (mm)
E
Bregma: -3.40 mm
F
antPAGdm
G
antPAGdl
H
antPAGl
cfos density (# cells/µm²)
Ctrl
ChR2
I
Bregma: -5.02 mm
J
posPAGdm
K
posPAGl
L
posPAGvl

**Supplementary figure 2, related to Figure 2 - The aVMHvl$^{PR+}$ and pVMHvl$^{PR+}$ subpopulations differ in their connectivity patterns to the PAG. Optogenetic activation of the aVMHvl$^{PR+}$ neurons did not induce changes in cFos density in the anterior and posterior portions of the PAG.** (A) Schematic of the quantified projections from pVMHvl$^{PR+}$ neurons to the PAG. (B) Representative image showing SynGFP expression in the PAG after an injection in the pVMHvl. Scale bar: 1000 µm (insect: scale bar 250 µm). (C) Heatmaps of binned SynGFP signal, after injections targeting aVMHvl$^{PR+}$ and pVMHvl$^{PR+}$ neurons, across different AP levels of the PAG, normalized to injection size and averaged across mice. Scale bar: 1000 µm. (D) Quantitative analysis of the normalized density of the SynGFP signal along the AP extent of the PAG. Mixed-effects test with repeated measures. Mean (±SD); *$p < 0.05$. Dark cyan: aVMHvl$^{PR+}$ projections, N = 5; Light cyan: pVMHvl$^{PR+}$ projections, N = 6. (F,I) Atlas slice containing the PAG levels considered as anterior and posterior PAG, respectively. (F-H, J-K) cFos density in the different subcolumns of the PAG along in two positions of its AP axis: (F-H) anterior, (J-K) posterior. Gray: Ctrl, N = 6, blue: ChR2, N = 8.

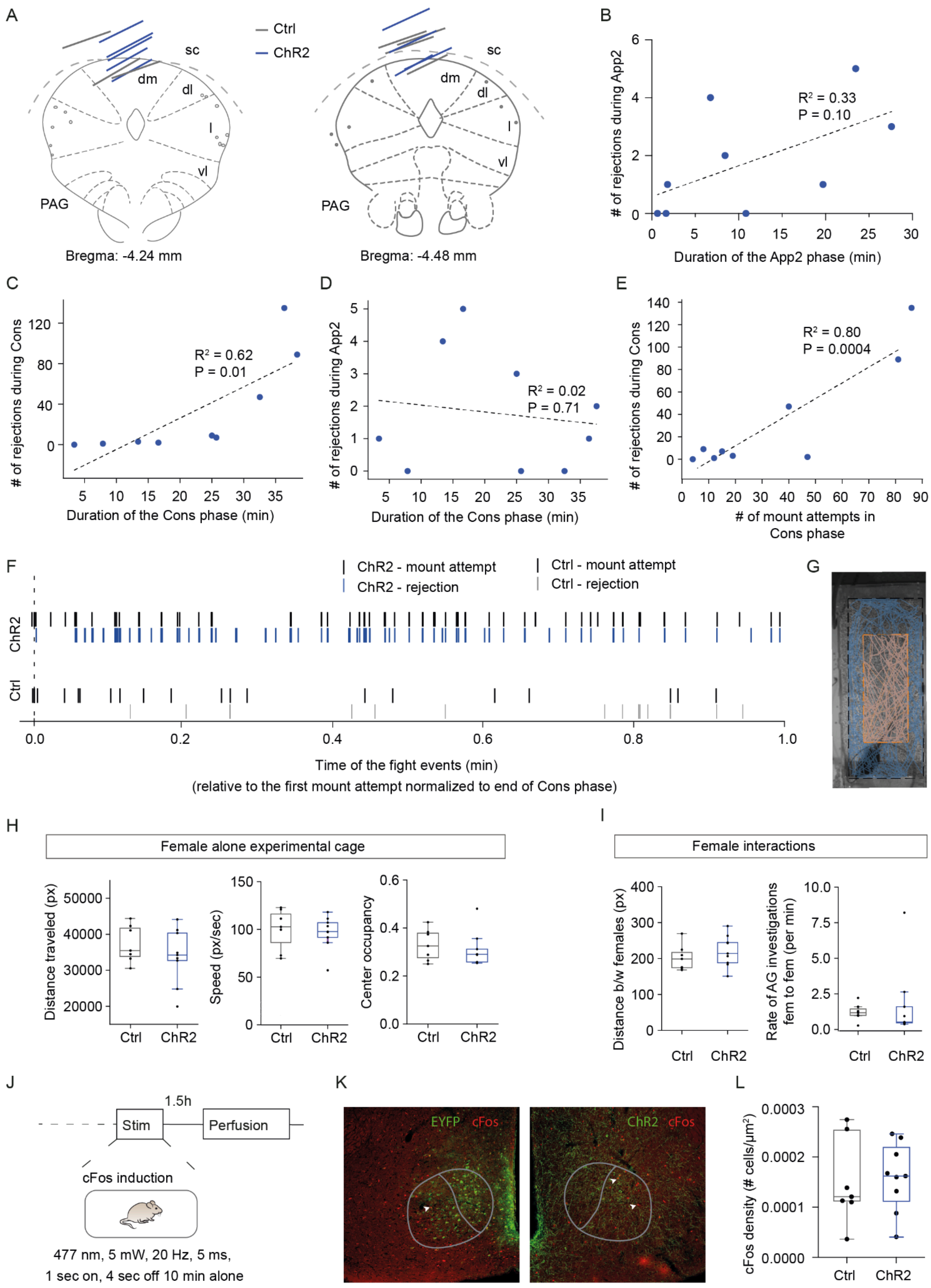

A
Ctrl
ChR2
sc
dm
dl
l
vl
PAG
Bregma: -4.24 mm
Bregma: -4.48 mm
B
R² = 0.33
P = 0.10
### of rejections during App2
Duration of the App2 phase (min)
C
R² = 0.62
P = 0.01
### of rejections during Cons
Duration of the Cons phase (min)
D
R² = 0.02
P = 0.71
### of rejections during App2
Duration of the Cons phase (min)
E
R² = 0.80
P = 0.0004
### of rejections during Cons
### of mount attempts in Cons phase
F
ChR2 - mount attempt
ChR2 - rejection
Ctrl - mount attempt
Ctrl - rejection
ChR2
Ctrl
Time of the fight events (min)
(relative to the first mount attempt normalized to end of Cons phase)
G
H
Female alone experimental cage
Distance traveled (px)
Speed (px/sec)
Center occupancy
Ctrl
ChR2
I
Female interactions
Distance b/w females (px)
Rate of AG investigations fem to fem (per min)
J
1.5h
Stim
Perfusion
cFos induction
477 nm, 5 mW, 20 Hz, 5 ms,
1 sec on, 4 sec off 10 min alone
K
EYFP
cFos
ChR2
cFos
L
cFos density (# cells/μm²)

**Supplementary Figure 3, related to Figure 3 - Effect of aVMHvl$^{PR+}$ to dmPAG afferents activation on rejection behavior and duration of the socio-sexual interaction phases. Optogenetic activation of aVMHvl$^{PR+}$ to dmPAG afferents does not lead to abnormalities in locomotion, anxiety or social cue investigation, and did not induce cFos increase in the aVMHvl.** (A) Schematic representation of the fiber tip locations (coloured lines) for ChR2 (blue) and Ctrl (grey) females, with each line representing a single mouse. (B) Rate of rejections during App2 as function of duration of App2 phase. (C) Rate of rejections during Cons as function of duration of Cons phase. (D) Rate of rejections during App2 as function of duration of Cons phase. (E) Rate of rejections during Cons as function of mount attempts during Cons phase. (F) Representative raster plot showing the occurrence of mount attempts displayed by the male and the corresponding sexual rejection displayed by one ChR2 female (top, black and blue) and one Ctrl female (bottom, black and grey) along the session (Appetitive 2 and Consummatory phases). Zero represents the first mount attempt. (G) Representative example of female's position tracked when alone in the experimental cage (blue+orange). The female´s occupancy of the center of the arena is represented in orange. (H) Distance traveled by the females when alone in the experimental cage, velocity of the females, and fraction of total time spent in the cage center when alone in the experimental cage. (I) Distance from an intruder stimulus female and rate of anogenital investigations towards the intruder female. During these sessions, no rejection (kicking and boxing) was observed. (J) Experimental design and timeline to quantify cFos expression after stimulation of aVMHvlPR+ axons at medial dmPAG. (K) Representative images showing cFos (red) and EYFP (left, green), or ChR2 (right, green), expression in the aVMHvl. White arrows highlight example cFos positive cells. Scale bar: 100 µm. (L) Number of cFos positive cells in the aVMHvl of EYFP and ChR2-expressing mice. Boxplots represent the median and interquartile range (IQR); whiskers extend to 1.53 IQR. Gray: Ctrl N = 7, blue: ChR2 N = 9.

**Table S1, related to Figure 1 and S1 - Output regions of aVMHvl$^{PR+}$ neurons.** Outputs, their abbreviations, their general group based on the Interactive Allen Brain Atlas classification (Red: Hypothalamus; Orange: Hindbrain; Violet: Midbrain; Salmon: Thalamus; Grey: Pallidum, Green: Cerebral Cortex, Blue: Striatum, Yellow: Cerebellum Pink: Medulla) and the density of Syn-GFP, in descending order.

| Brain Regions | Abbreviation | Average of the Synaptophysin-GFP density, normalized to injection size |
|---|---|---|
| Anterior hypothalamic nucleus | AHN | 9.26214 |
| Periaqueductal gray | PAG | 7.54088 |
| Retrochiasmatic area | RCH | 7.36219 |
| Accessory supraoptic group | ASO | 6.86441 |
| Subparaventricular zone | SBPV | 6.84550 |
| Precommissural nucleus | PRC | 5.55556 |
| Ventromedial preoptic nucleus | VMPO | 4.90620 |
| Medial preoptic nucleus | MPN | 4.15519 |
| Nucleus of the posterior commissure | NPC | 4.11978 |
| Peripeduncular nucleus | PP | 3.66420 |
| Posterior hypothalamic nucleus | PH | 3.57576 |
| Supraoptic nucleus | SO | 3.53575 |
| Suprachiasmatic nucleus | SCH | 3.39911 |
| Medial preoptic area | MPOA | 3.39763 |
| Ventral premammillary nucleus | PMv | 3.37544 |
| Tuberal nucleus | TU | 3.30646 |
| Dorsal premammillary nucleus | PMd | 3.23551 |
| Bed nuclei of the stria terminalis | BNST | 3.17918 |
| Posterodorsal preoptic nucleus | PD | 3.13801 |
| Paraventricular hypothalamic nucleus | PVH | 3.13356 |
| Subparafascicular area | SPF | 3.07428 |
| Dorsomedial nucleus of the hypothalamus | DMH | 2.97908 |
| Paraventricular nucleus of the thalamus | PVT | 2.92383 |
| Lateral terminal nucleus of the accessory optic tract | LT | 2.69898 |
| Posterior intralaminar thalamic nucleus | PIL | 2.61474 |
| Perifornical nucleus | PeF | 2.44879 |
| Intermediate geniculate nucleus | IntG | 2.31123 |
| Parataenial nucleus | PT | 2.30966 |

| | | |
|---|---|---|
| Supramammillary nucleus | SUM | 2.26364 |
| Parastrial nucleus | PS | 2.16233 |
| Dorsal nucleus raphe | DR | 2.14467 |
| Ventrolateral preoptic nucleus | VLPO | 2.03960 |
| Arcuate hypothalamic nucleus | ARH | 2.00383 |
| Tuberomammillary nucleus | TM | 1.97822 |
| Superior colliculus | SC | 1.97174 |
| Nucleus of reuniens | RE | 1.95290 |
| Anterodorsal preoptic nucleus | ADP | 1.85357 |
| Subparafascicular nucleus | SPF | 1.69693 |
| Anteroventral preoptic nucleus | AVP | 1.59468 |
| Cuneiform nucleus | CUN | 1.54590 |
| Lateral hypothalamic area | LHA | 1.45503 |
| Anteroventral periventricular nucleus | AVPV | 1.42618 |
| Lateral preoptic area | LPO | 1.38791 |
| Midbrain reticular nucleus | MRN | 1.22753 |
| Interanteromedial nucleus of the thalamus | IAM | 1.21258 |
| Posterior pretectal nucleus | PPT | 1.20975 |
| Periventricular hypothalamic nucleus | PVa | 1.13762 |
| Median eminence | ME | 1.08594 |
| Xiphoid thalamic nucleus | Xi | 1.04850 |
| Posterior limiting nucleus of the thalamus | POL | 0.87664 |
| Parasubthalamic nucleus | PSTN | 0.87442 |
| Median preoptic nucleus | MEPO | 0.80622 |
| Intergeniculate leaflet of the lateral geniculate complex | IGL | 0.78470 |
| Nucleus sagulum | SAG | 0.77381 |
| Interfascicular nucleus raphe | IF | 0.74984 |
| Medial septal nucleus | MS | 0.74601 |
| Zona incerta | ZI | 0.72150 |
| Parabrachial nucleus | PB | 0.70698 |
| Rostral linear nucleus raphe | RL | 0.66935 |
| Perireunensis nucleus | PR | 0.66592 |
| Pedunculopontine nucleus | PPN | 0.65948 |
| Interanterodorsal nucleus of the thalamus | IAD | 0.65620 |
| Central linear nucleus raphe | CLI | 0.64680 |

| | | |
|---|---|---|
| Tegmental reticular nucleus | TRN | 0.64476 |
| Central medial nucleus of the thalamus | CM | 0.63851 |
| Secondary motor area | MOs | 0.63765 |
| Ventral tegmental area | VTA | 0.62737 |
| Lateral septal nucleus | LS | 0.62242 |
| Laterodorsal tegmental nucleus | LDT | 0.51920 |
| Central amygdalar nucleus | CEA | 0.51845 |
| Pallidum | PAL | 0.50468 |
| Lateral amygdalar nucleus | LA | 0.50149 |
| Rhomboid nucleus | RH | 0.49069 |
| Pontine gray | PG | 0.47021 |
| Diagonal band nucleus | NBD | 0.46538 |
| Perirhinal area | PERI | 0.46481 |
| Striatum | STR | 0.46417 |
| Infralimbic area | ILA | 0.46260 |
| Intermediodorsal nucleus of the thalamus | IMD | 0.45022 |
| Lobule II | CENT2 | 0.44999 |
| Pontine central gray | PCG | 0.44632 |
| Pontine reticular nucleus | PRNc | 0.44196 |
| Lateral habenula | LH | 0.44160 |
| Ectorhinal area/Layer 5 | ECT5 | 0.43864 |
| Lateral mammillary nucleus | LM | 0.43605 |
| Medial mammillary nucleus | MM | 0.43132 |
| Retroparafascicular nucleus | RPF | 0.43072 |
| Anterior cingulate area | ACA | 0.42741 |
| Orbital area | ORB | 0.42583 |
| Substantia innominata | SI | 0.39700 |
| Basolateral amygdalar nucleus | BLA | 0.38962 |
| Fields of Forel | FF | 0.38841 |
| Red nucleus | RN | 0.38495 |
| Medial amygdalar nucleus | MEA | 0.38038 |
| Cortical amygdalar area | COA | 0.36299 |
| Entorhinal area | ENT | 0.35750 |
| Central lateral nucleus of the thalamus | CL | 0.33396 |
| Prosubiculum | ProS | 0.33264 |
| Basomedial amygdalar nucleus | BMA | 0.33159 |
| Primary somatosensory area | SSp | 0.33138 |

| | | |
|---|---|---|
| Ventral part of the lateral geniculate complex | LGv | 0.33078 |
| Interstitial nucleus of Cajal | INC | 0.32825 |
| Posterior amygdalar nucleus | PA | 0.32163 |
| Pons | P | 0.31685 |
| Supratrigeminal nucleus | SUT | 0.31391 |
| Intercalated amygdalar nucleus | IA | 0.31091 |
| Agranular insular area | AI | 0.30961 |
| Postrhinal area | VISpor | 0.30665 |
| Suprageniculate nucleus | SGN | 0.30287 |
| Superior central nucleus raphe | CS | 0.30248 |
| Ectorhinal area/Layer 6a | ECT6a | 0.30062 |
| Claustrum | CLA | 0.30058 |
| Medial geniculate complex | MG | 0.27160 |
| Nucleus of the lateral lemniscus | NLL | 0.27091 |
| Endopiriform nucleus | EP | 0.26136 |
| Cortical subplate | CTXsp | 0.26093 |
| Nucleus raphe pontis | RPO | 0.24859 |
| Anterior pretectal nucleus | APN | 0.24645 |
| Primary motor area | MOp | 0.24353 |
| Superior olivary complex | SOC | 0.24315 |
| Principal sensory nucleus of the trigeminal | PSV | 0.22654 |
| Lateral posterior nucleus of the thalamus | LP | 0.22620 |
| Mediodorsal nucleus of thalamus | MD | 0.22509 |
| Posterior triangular thalamic nucleus | PoT | 0.20893 |
| Subthalamic nucleus | STN | 0.20222 |
| Prelimbic area | PL | 0.20073 |
| Reticular nucleus of the thalamus | RT | 0.19039 |
| Nucleus of the optic tract | NOT | 0.18173 |
| Ectorhinal area/Layer 2/3 | ECT2/3 | 0.17575 |
| Inferior colliculus | IC | 0.17387 |
| Substantia nigra | SNr | 0.17200 |
| Anterior amygdalar area | AAA | 0.17042 |
| Piriform-amygdalar area | PAA | 0.16909 |
| Temporal association areas | TEa | 0.16055 |
| Subiculum | SUB | 0.15837 |
| Globus pallidus | GPe | 0.15658 |

| | | |
|---|---|---|
| Dorsal part of the lateral geniculate complex | LGd | 0.14549 |
| Peritrigeminal zone | P5 | 0.13732 |
| Anteromedial nucleus | AM | 0.12585 |
| Ventral medial nucleus of the thalamus | VM | 0.12458 |
| Frontal pole. layer 5 | FRP5 | 0.12376 |
| Postpiriform transition area | TR | 0.11408 |
| Lobules IV-V | CUL4.5 | 0.11363 |
| Nucleus accumbens | ACB | 0.11314 |
| Field CA3 | CA3 | 0.10620 |
| Visceral area | VISC | 0.10592 |
| Caudoputamen | CP | 0.10153 |
| Field CA1 | CA1 | 0.10135 |
| Lateral dorsal nucleus of thalamus | LD | 0.10005 |
| Postsubiculum | POST | 0.09834 |
| Anteroventral nucleus of thalamus | AV | 0.09772 |
| Retrosplenial area | RSP | 0.09290 |
| Piriform area | PIR | 0.07776 |
| Primary visual area | VISp | 0.07204 |
| Simple lobule | SIM | 0.05893 |
| Ventral posterolateral nucleus of the thalamus | VPLpc | 0.05670 |
| Ventral anterior-lateral complex of the thalamus | VAL | 0.04957 |
| Olfactory tubercle | OT | 0.04842 |
| Dentate gyrus | DG | 0.04692 |
| Anterior olfactory nucleus | AON | 0.02049 |
| Supplemental somatosensory area | SSs | 0.01469 |

**Table S2, related to Figure S1 - Output regions of pVMHvl$^{PR+}$ neurons.** Outputs, their abbreviations, their general group based on the Interactive Allen Brain Atlas classification and the density of Syn-GFP, in a descending order.

| Brain Regions | Abbreviation | Average of the Synaptophysin-GFP density, normalized to injection size |
|---|---|---|
| Posterodorsal preoptic nucleus | PD | 36.97323 |
| Medial preoptic nucleus | MPN | 21.88729 |
| Barrington's nucleus | B | 16.24492 |
| Retrochiasmatic area | RCH | 13.08515 |
| Medial preoptic area | MPOA | 12.48399 |
| Anterior hypothalamic nucleus | AHN | 11.82536 |
| Ventromedial preoptic nucleus | VMPO | 10.53193 |
| Ventral premammillary nucleus | PMv | 10.29004 |
| Anterodorsal preoptic nucleus | ADP | 9.88758 |
| Periaqueductal gray | PAG | 9.09563 |
| Tuberal nucleus | TU | 8.53331 |
| Perifornical nucleus | PeF | 8.01487 |
| Subparaventricular zone | SBPV | 7.81557 |
| Parastrial nucleus | PS | 7.73915 |
| Paraventricular hypothalamic nucleus | PVH | 7.70773 |
| Anteroventral preoptic nucleus | AVP | 7.66665 |
| Anteroventral periventricular nucleus | AVPV | 6.94925 |
| Paranigral nucleus | PN | 6.61468 |
| Locus ceruleus | LC | 6.56365 |
| Accessory supraoptic group | ASO | 6.48413 |
| Supramammillary nucleus | SUM | 5.57781 |
| Tuberomammillary nucleus | TM | 5.39667 |
| Bed nuclei of the stria terminalis | BNST | 5.23984 |
| Arcuate hypothalamic nucleus | ARH | 5.17939 |
| Midbrain trigeminal nucleus | MEV | 5.07196 |
| Dorsal premammillary nucleus | PMd | 5.01258 |
| Lateral preoptic area | LPO | 4.82732 |
| Posterior hypothalamic nucleus | PH | 4.68860 |
| Dorsal nucleus raphe | DR | 4.40766 |
| Suprachiasmatic nucleus | SCH | 4.28694 |
| Ventrolateral preoptic nucleus | VLPO | 4.16274 |
| Subparafascicular area | SPF | 4.11067 |
| Interfascicular nucleus raphe | IF | 3.99609 |
| Median preoptic nucleus | MEPO | 3.67428 |

| | | |
|---|---|---|
| Central linear nucleus raphe | CLI | 3.60864 |
| Lateral hypothalamic area | LHA | 3.46585 |
| Periventricular hypothalamic nucleus | PVa | 3.41054 |
| Paraventricular nucleus of the thalamus | PVT | 3.27068 |
| Ventral tegmental area | VTA | 2.98361 |
| Posterior intralaminar thalamic nucleus | PIL | 2.89854 |
| Dorsomedial nucleus of the hypothalamus | DMH | 2.75169 |
| Precommissural nucleus | PRC | 2.52601 |
| Cuneiform nucleus | CUN | 2.39953 |
| Sublaterodorsal nucleus | SLD | 2.39387 |
| Parataenial nucleus | PT | 2.31468 |
| Nucleus of the posterior commissure | NPD | 2.24238 |
| Subceruleus nucleus | SLC | 2.20964 |
| Pontine central gray | PCG | 2.11195 |
| Subparafascicular nucleus | SPF | 2.08222 |
| Parasubthalamic nucleus | PSTN | 2.04378 |
| Midbrain reticular nucleus | RR | 2.03495 |
| Parabrachial nucleus | PB | 2.02998 |
| Edinger-Westphal nucleus | EW | 2.00957 |
| Laterodorsal tegmental nucleus | LDT | 1.94212 |
| Lateral mammillary nucleus | LM | 1.92730 |
| Peripeduncular nucleus | PP | 1.92290 |
| Anterior tegmental nucleus | AT | 1.90825 |
| Supraoptic nucleus | SO | 1.85126 |
| Pedunculopontine nucleus | PPN | 1.82956 |
| Nucleus raphe pontis | RPO | 1.74247 |
| Medial septal nucleus | MS | 1.59838 |
| Nucleus of reuniens | RE | 1.56716 |
| Lateral septal nucleus | LS | 1.54146 |
| Median eminence | ME | 1.38409 |
| Superior central nucleus raphe | CS | 1.35726 |
| Medial terminal nucleus of the accessory optic tract | MT | 1.34157 |
| Lateral terminal nucleus of the accessory optic tract | LT | 1.19808 |
| Medial mammillary nucleus | MM | 1.17073 |
| Diagonal band nucleus | NDB | 1.12962 |

| | | |
|---|---|---|
| Frontal pole. layer 5 | FRP5 | 1.11785 |
| Rostral linear nucleus raphe | RL | 1.09303 |
| Xiphoid thalamic nucleus | Xi | 1.06203 |
| Ventral tegmental nucleus | VTN | 1.02180 |
| Interpeduncular nucleus | IPN | 0.92716 |
| Supratrigeminal nucleus | SUT | 0.91355 |
| Nucleus incertus | NI | 0.91120 |
| Nucleus raphe magnus | RM | 0.87743 |
| Striatum | Striatum | 0.84194 |
| Posterior amygdalar nucleus | PA | 0.84032 |
| Retroparafascicular nucleus | RPF | 0.82734 |
| Intermediodorsal nucleus of the thalamus | IMD | 0.78421 |
| Perireunensis nucleus | PR | 0.71721 |
| Lateral habenula | LH | 0.71502 |
| Red nucleus | RN | 0.69788 |
| Zona incerta | ZI | 0.69037 |
| Pallidum | Pallidum | 0.66607 |
| Nucleus of Darkschewitsch | ND | 0.65215 |
| Substantia innominata | SI | 0.64746 |
| Interstitial nucleus of Cajal | INC | 0.64620 |
| Pontine reticular nucleus | PRNc | 0.63333 |
| Central amygdalar nucleus | CEA | 0.61972 |
| Superior colliculus | SCs | 0.60829 |
| Magnocellular reticular nucleus | MARN | 0.60573 |
| Posterior limiting nucleus of the thalamus | POL | 0.57866 |
| Peritrigeminal zone | P5 | 0.54558 |
| Fields of Forel | FF | 0.53212 |
| Pons | P | 0.51967 |
| Orbital area | ORB | 0.46207 |
| Interanterodorsal nucleus of the thalamus | IAD | 0.41671 |
| Medial amygdalar nucleus | MEA | 0.41175 |
| Nucleus sagulum | SAG | 0.41080 |
| Interanteromedial nucleus of the thalamus | IAM | 0.40043 |
| Subthalamic nucleus | STN | 0.39385 |
| Mediodorsal nucleus of thalamus | MD | 0.38785 |
| Tegmental reticular nucleus | TRN | 0.35724 |

| | | |
|---|---|---|
| Posterior pretectal nucleus | PPT | 0.34989 |
| Central medial nucleus of the thalamus | CM | 0.34721 |
| Prelimbic area | PL | 0.32755 |
| Substantia nigra | SNr | 0.28491 |
| Secondary motor area | MOs | 0.28216 |
| Nucleus accumbens | ACB | 0.27082 |
| Basomedial amygdalar nucleus | BMA | 0.26182 |
| Ventral posteromedial nucleus of the thalamus | VPM | 0.25333 |
| Intercalated amygdalar nucleus | IA | 0.23851 |
| Posterior triangular thalamic nucleus | PoT | 0.23440 |
| Anterior cingulate area | ACA | 0.23359 |
| Anterior pretectal nucleus | APN | 0.23351 |
| Gigantocellular reticular nucleus | GRN | 0.23224 |
| Nucleus of the brachium of the inferior colliculus | NB | 0.23048 |
| Reticular nucleus of the thalamus | RT | 0.22872 |
| Postsubiculum | POST | 0.22643 |
| Medial vestibular nucleus | MV | 0.22369 |
| Taenia tecta | TT | 0.21130 |
| Superior olivary complex | SOC | 0.20049 |
| Hippocampo-amygdalar transition area | HATA | 0.19888 |
| Medial geniculate complex | MG | 0.19886 |
| Paragigantocellular reticular nucleus | PGRN | 0.19695 |
| Parafascicular nucleus | PF | 0.19582 |
| Hippocampal formation | HPF | 0.19371 |
| Septofimbrial nucleus | SF | 0.18806 |
| Presubiculum | PRE | 0.18795 |
| Infralimbic area | ILA | 0.18714 |
| Cortical amygdalar area | COA | 0.16860 |
| Inferior olivary complex | IO | 0.16645 |
| Dorsal peduncular area | DP | 0.16235 |
| Agranular insular area | AI | 0.15711 |
| Cortical subplate | CTXsp | 0.15381 |
| Anterior amygdalar area | AAA | 0.15152 |
| Ventral medial nucleus of the thalamus | VM | 0.15102 |
| Magnocellular nucleus | MA | 0.14124 |
| Parvicellular reticular nucleus | PARN | 0.13624 |

| | | |
|---|---|---|
| Intermediate reticular nucleus | IRN | 0.13447 |
| Koelliker-Fuse subnucleus | KF | 0.12984 |
| Retrosplenial area | RSP | 0.12977 |
| Facial motor nucleus | VII | 0.12791 |
| Lobule II | CENT2 | 0.12217 |
| Lateral reticular nucleus | LRN | 0.11786 |
| Basolateral amygdalar nucleus | BLA | 0.11666 |
| Motor nucleus of trigeminal | V | 0.11523 |
| Central lateral nucleus of the thalamus | CL | 0.11446 |
| Primary motor area | Mop | 0.11392 |
| Nucleus of the solitary tract | NTS | 0.10746 |
| Subiculum | SUB | 0.10044 |
| Anterior olfactory nucleus | AON | 0.09810 |
| Olfactory areas | OLF | 0.09299 |
| Parasubiculum | PAR | 0.09113 |
| Inferior colliculus | IC | 0.08747 |
| Ventral anterior-lateral complex of the thalamus | VAL | 0.08180 |
| Principal sensory nucleus of the trigeminal | PSV | 0.07901 |
| Nucleus prepositus | PRP | 0.07032 |
| Triangular nucleus of septum | TRS | 0.06953 |
| Prosubiculum | ProS | 0.06862 |
| Ventral part of the lateral geniculate complex | LGv | 0.06667 |
| Lobule III | CENT3 | 0.06567 |
| Superior vestibular nucleus | SUV | 0.06113 |
| Pontine gray | PG | 0.05881 |
| Lateral amygdalar nucleus | LA | 0.05785 |
| Nucleus of the lateral lemniscus | NLL | 0.05650 |
| Field CA3 | CA3 | 0.05183 |
| Anteroventral nucleus of thalamus | AV | 0.05031 |
| Field CA1 | CA1 | 0.04712 |
| Entorhinal area | ENT | 0.04677 |
| Globus pallidus | GP | 0.04472 |
| Lateral posterior nucleus of the thalamus | LP | 0.04445 |
| Endopiriform nucleus | EP | 0.04398 |
| Spinal nucleus of the trigeminal | SPVC | 0.04326 |
| Lateral dorsal nucleus of thalamus | LD | 0.04316 |

| | | |
|---|---|---|
| Main olfactory bulb | MOB | 0.04282 |
| Lobules IV-V | CUL4.5 | 0.04260 |
| Olfactory tubercle | OT | 0.03773 |
| Spinal vestibular nucleus | SPIV | 0.03624 |
| Nodulus (X) | NOD | 0.03399 |
| Caudoputamen | CP | 0.03386 |
| Piriform-amygdalar area | PAA | 0.03214 |
| Postpiriform transition area | TR | 0.03209 |
| Dentate gyrus | DG | 0.02450 |
| Piriform area | PIR | 0.01976 |
| Primary visual area | VISp | 0.01947 |
| Posterior complex of the thalamus | PO | 0.01681 |
| Flocculus | FL | 0.01210 |
| Simple lobule | SIM | 0.01150 |
| Paramedian lobule | PRM | 0.01037 |
| Crus 1 | ANcr1 | 0.00633 |
| Paraflocculus | PFL | 0.00381 |
